# Age-related and heteroplasmy-related variation in human mtDNA copy number

**DOI:** 10.1101/030205

**Authors:** Manja Wachsmuth, Alexander Hübner, Mingkun Li, Burkhard Madea, Mark Stoneking

**Author notes:** Equal contribution (Manja Wachsmuth, Alexander Hübner).

## Abstract

The mitochondrial (mt) genome is present in many copies in human cells, and intra-individual variation in mtDNA sequences is known as heteroplasmy. Recent studies found that heteroplasmies are highly tissue-specific, site-specific, and allele-specific, however the functional implications have not been explored. This study investigates variation in mtDNA copy numbers (mtCN) in 12 different tissues obtained at autopsy from 152 individuals (ranging in age from 3 days to 96 years). Three different methods to estimate mtCN were compared: shotgun sequencing (in 4 tissues), capture-enriched sequencing (in 12 tissues) and droplet digital PCR (ddPCR, in 2 tissues). The highest precision in mtCN estimation was achieved using shotgun sequencing data. However, capture-enrichment data provide reliable estimates of relative (albeit not absolute) mtCNs. Comparisons of mtCN from different tissues of the same individual revealed that mtCNs in different tissues are, with few exceptions, uncorrelated. Hence, each tissue of an individual seems to regulate mtCN in a tissue-related rather than an individual-dependent manner. Skeletal muscle (SM) samples showed an age-related decrease in mtCN that was especially pronounced in males, while there was an age-related increase in mtCN for liver (LIV) samples. MtCN in SM samples was significantly negatively correlated with both the total number of heteroplasmic sites and with minor allele frequency (MAF) at two heteroplasmic sites, 408 and 16327. Heteroplasmies at both sites are highly specific for SM, accumulate with aging and are part of functional elements that regulate mtDNA replication. These data support the hypothesis that selection acting on these heteroplasmic sites is reducing mtCN in SM of older individuals.

**Author Summary:** The total number of mitochondrial genomes in a human cell differs between individuals and between the tissues of a single individual; however the factors that influence this variation remain unknown. We estimated mtDNA copy number (mtCN) in 12 different tissues of 152 individuals applying three different methods, and found age-related variation for two tissues: mtCN in skeletal muscle is negatively correlated with age (especially in males) while mtCN in liver is positively correlated with age. Overall, mtCNs of different tissues within an individual are mainly independent of each other, indicating that tissue-specific rather than individual-specific processes largely influence mtCN.

Heteroplasmy refers to intra-individual differences in the sequence of the mtDNA genome and heteroplasmic mutations accumulate during aging. Linear and partial regression analyses of mtCN with heteroplasmy (determined in a previous study of these same samples) revealed that the decrease of mtCN in skeletal muscle is mainly correlated with an increasing total number of heteroplasmic sites, and with increasing minor allele frequency at two sites (408 and 16327), that are heteroplasmic almost exclusively in skeletal muscle. As both sites are part of functional elements required for regulation of mtDNA replication, we suggest that selection may be acting via increasing heteroplasmy to reduce mtCN during aging.

## Introduction

Mitochondria are the central cellular structures for energy production. While most human mt proteins are encoded in the nuclear genome and imported to the mitochondrion following translation, mitochondria also contain their own genome [1]. In addition to tRNA and rRNA genes, the mt genome harbors 13 genes encoding proteins of the respiratory chain. The human mt genome is usually present in several copies per mitochondrion and the total number of mtDNA copies per cell varies greatly between different individuals and different tissues of the same individual [2, 3]. For example, myocardial muscle cells contain on average more than 6,000 copies per cell [4], while leukocytes have around 350 mtDNA copies per cell [5].

Replication and degradation of mtDNA are coupled in order to keep mtDNA levels constant in a cell [3]. Changes in mtCN have been correlated with several diseases and some factors that influence mtCN have been described [3, 6]. These are for example replication regulating proteins such as the single-stranded-DNA-binding protein mtSSB, which stimulates mitochondrial DNA polymerase γ and therefore increases replication rate [6]. In addition, overexpression of mitochondrial RNA polymerase and its mitochondrial transcription factor A (TFAM) enhance replication by increased synthesis of primers for replication [6]. A reduction of TFAM in heterozygous knockout mice showed a strong reduction in mtCN and a total knockout resulted in a total lack of mtDNA and lethality [7].

However, it is still not known how inter-individual differences arise in mtCN, to what extent such inter-individual differences in mtCN are correlated across different tissues of an individual, and what effect these natural differences might have on healthy individuals. Several studies have examined changes in mtCN with age; while some detected a decrease of copy number with increasing age in healthy individuals [8-10], others failed to identify age-dependent changes in copy number [4, 11] or showed an increase for certain tissues [12]. Thus, there is currently much uncertainty concerning the role of age and other factors on mtCN.

Copy number is usually estimated by quantitative PCR methods [4, 10], although some studies have utilized shotgun sequencing data [9, 13-15]. However, comparisons of different methods for estimating mtCN are rarely done [9] and moreover most studies have focused on a single tissue. Here, we analyze mtCN in 12 different tissues that were obtained at autopsy from 152 individuals. We compared three different methods for estimating mtCN: ddPCR (in 2 tissues); shotgun sequencing (in 4 tissues); and capture-enriched sequencing (in all 12 tissues). Furthermore, we examined the influence of various features such as age, haplogroup and sex on mtCN.

We also inquired whether mtCN can be linked to mtDNA heteroplasmy, i.e. intra-individual variability in the mtDNA sequence. Several studies revealed that heteroplasmy increases in healthy human individuals with aging [16-19]. In addition, it has been shown that there is a tissue-specific accumulation of heteroplasmy at defined sites [15, 17-21], which might hint to either positive selection on specific sites [15, 19] or a lack of negative selection due to relaxation of functional constraints [22]. All samples examined for mtCN in the present study were investigated previously for heteroplasmy [19]. As the most common heteroplasmic sites are found exclusively in the control region of the mitochondrial genome, which is essential for replication and transcription initiation and regulation [23, 24], we hypothesized that heteroplasmic mutations in the control region could have an influence on mtCN, and indeed we report here that an increase in the minor allele frequency (MAF) at two sites that are frequently heteroplasmic in skeletal muscle is associated with an age-related decrease in mtCN.

## Materials and Methods

### Tissue collection and DNA extraction

Twelve different tissues (blood: BL; cerebellum: CEL; cerebrum: CER; cortex: CO; kidney: KI; large intestine: LI; liver: LIV myocardial muscle: MM; ovary: OV; small intestine: SI; skin: SK; and skeletal muscle: SM) were obtained at autopsy from 152 human individuals (85 males, 67 females) of mostly European origin and DNA was extracted as previously described [19]. The collection of samples and the experimental procedures were approved by the Ethics Commissions of the Rheinische Friedrich Wilhelm University Medical Faculty and of the University of Leipzig Medical Faculty (Approval numbers: Rheinische Friedrich Wilhelm University: 121/11, University of Leipzig: 349-11-07112011 and 305-15-24082015).

### MtDNA quantification using shotgun sequencing data and capture enriched data

Libraries for sequencing were prepared as previously described [19]. Shotgun sequencing (without mtDNA enrichment) was performed on an Illumina HiSeq platform using 95 base pair, paired-end reads. Two runs were performed: run 1 consisted of pooled libraries from four tissues (with all samples from BL (148 samples), SM (152), LIV (152) and MM (150)) and run 2 consisted of the same libraries from two tissues, BL and SM. To check reproducibility of the results with independent libraries, we prepared a new subset of libraries from the BL and SM DNA extracts from each of 26 individuals of mixed age, sex and mtCN and performed shotgun sequencing on an Illumina MiSeq platform using 150 base pair, paired-end reads (referred to as shotgun sequencing run 3).

Reads of at least 50 bp length were mapped against the human reference genome 19 (hg19) using BWA with default settings [25] and without filtering for high mapping quality. All reads that aligned successfully, i.e. both mates of a pair mapped to the same chromosome and within a distance to each other of at most 3 standard deviations of the insert size distribution, were analyzed and the length of their insert sizes summed up per chromosome. For each chromosome the number of mappable bases was determined by removing any poly-N stretch in the hg19 (> 5 consecutive Ns) and counting the number of remaining bases. The number of aligned bases of each chromosome was divided by the number of mappable bases on this chromosome to obtain the coverage per chromosome and sample.

Samples were subsequently removed from the data set when either of the following criteria was fulfilled: 1) the number of bases aligning to mtDNA was ≤ 60% of the total mtDNA length; 2) the logarithmic normalized SD of the autosomal coverage was identified by a Grubbs test [26] as an outlier.

For each tissue the standard deviation of the coverage of each autosome was determined over all remaining samples. Chromosomes that were identified as an outlier by a Grubbs test based on their SD in 50% of the tissues were not considered in the mtCN calculation. The mtCN was determined by the following formula [9]:

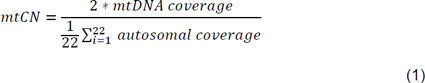

In order to account for the difference in absolute mtCN per tissue, the SDs were normalized by the mean autosomal coverage of a sample.

In a previous study, Illumina sequencing had been performed after capture-enrichment for mtDNA [19]. The capture-enriched sequencing data from samples from BL (139), CEL (150), CER (143), CO (152), KI (151), LI (150), LIV (151), MM (149), OV (47), SI (150), SK (152) and SM (150), were processed and mtCN estimation carried out as described above for shotgun sequencing data.

### MtDNA quantification using ddPCR

The mtCN from BL (150 samples) and SM (152) samples was determined using droplet digital PCR (ddPCR), a quantitative PCR method that allows an absolute measurement of the number of target DNA molecules [27]. A 20 µl PCR mix was prepared and dispersed into up to 20,000 droplets. After amplification, droplets that included template DNA were identified by fluorescent dyes and counted. Specific primers (S1 Table) for amplification of mt and nDNA regions were modified according to [10]. For determination of mtCN in BL, mtDNA and nDNA were amplified in the same multiplex reaction using HEX- and FAM-labeled probes for n and mtDNA, respectively. Reactions were performed in ddPCR Supermix including primers, probes and 1 ng template according to manufacturer’s instructions. Due to the high mtCN in SM, determination of nDNA and mtDNA in SM samples was performed in two separate reactions with 10 ng template for nDNA and 20 pg template for mtDNA and either nuclear or mt specific primers in EvaGreen Supermix. To control for differences between DNA preparations, at least 12 DNA samples per tissue type were extracted twice and the mtCN of different preparations was compared. MtCN was calculated after correction for dilution factors by

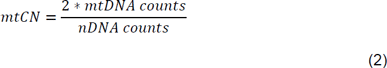

For each sample the arithmetic mean and the standard deviation (SD) of mtCN was calculated over all replicates. To account for the separate ddPCR reactions for nDNA and mtDNA measurement per SM sample, the SDs were averaged over the individual SDs of the SM samples. To compare the SDs between BL samples and SM samples, the SD was normalized by the mean mtCN of each sample. Both the arithmetic mean and the SD were subsequently averaged over all samples of a tissue type.

### Estimating the likelihood of detecting NUMTs

In order to estimate the likelihood that recent mtDNA insertions into the nuclear genome (NUMTs) could artificially increase the mtCN estimates from sequencing data, we analyzed the length distribution of identified NUMTs from a NUMT database [28] and compared it to the insert size distribution in our data set. When paired-end reads overlapped, the length of the merged read was defined as the insert size [29]. If reads were too far apart to be merged, the length of the region flanked by the two reads was defined as insert size. A read from a NUMT can only be aligned falsely with high quality to mtDNA when the read does not contain a flanking nuclear DNA signature of at least a few high-quality bases. For each insert size, all NUMTs longer than the insert were extracted from the published list as those inserts could potentially fully fall in a NUMT region. The probability of false alignment to the human mtDNA sequence was then estimated, taking into account the number of putative read insert positions within a NUMT, the number of NUMTs in the human genome that were larger than each read insert size, and the average coverage of all autosomes in each sequencing run, as follows:

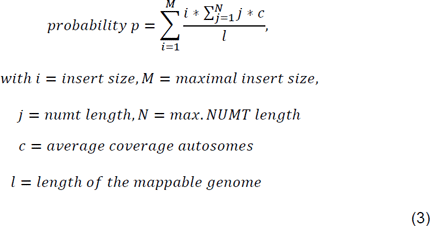

### Correlation of mtCN with individual parameters and with heteroplasmy

The mtCN values for each tissue were log-transformed to produce a normal distribution of the residuals for comparison with other parameters, e.g. age, sex, haplogroup, etc. An underlying normal distribution of the residuals is a requirement for many statistical tests (including the linear regression models used here) and was therefore formally tested for every tissue using a Shapiro-Wilk test. In order to fulfill the requirement of a normal distribution, one LIV-sample (LIV253) with very high mtCN had to be excluded from the shotgun sequencing data sets prior to further investigations. Linear regression and Pearson correlation analysis as implemented in R [30] was performed and the resulting p-values were corrected for multiple testing using the Benjamini-Hochberg method [31]. Partial regression analysis was used to investigate the combined effect of age and heteroplasmy levels on mtCN [32]. When performing linear regression analysis between MAF of specific heteroplasmies and mtCN, only heteroplasmies that were present in at least 10 individuals were considered. We tested for correlations of mtCN with single heteroplasmic sites, pairs of sites occurring in the same individual, and triplets of heteroplasmic sites, using linear regression and Pearson correlation analysis as described above.

## Results

### Reproducibility of mtCN estimation

For all analyses, tissue samples that had been collected at autopsy from 152 individuals and 12 different tissues in a previous study [19] were used. MtCN was estimated using shotgun sequencing (in BL, MM, LIV and SM), capture-enrichment sequencing (in BL, CEL, CER, CO, KI, LI, LIV, MM, OV, SI, SK and SM), and ddPCR (in BL and SM). The first two approaches are based on sequencing data and mtCN is estimated by dividing the number of bases that map to the mtDNA by the number of bases mapped to the nuclear genome. In order to estimate reproducibility, variation in coverage across the different autosomes was measured and chromosomes with high variation (as identified by the Grubbs outlier test) were removed. Chromosomes 16 and 19 were excluded from the mtCN calculation in the shotgun sequencing data as the SD of their coverage was enhanced in at least 50 % of the tissues according to the Grubbs test (S1 Fig). In addition, for all 12 tissues from all individuals, mtCNs were estimated using capture-enriched sequencing data. Here, sequencing libraries had been enriched for mtDNA prior to sequencing [19], and so mtCN estimates will be elevated in the capture-enriched data; however, relative mtCN estimates might be the same, and so allow for comparisons between individuals. According to the Grubbs outlier test, SD values for the coverage of chromosomes 1, 16 and 19 were enhanced and therefore these chromosomes were not included in mtCN calculations from capture-enriched sequences (S1 Fig).

MtDNA insertions into the nuclear genome (NUMTs) could potentially artificially increase mtCNs calculated from sequencing data if reads deriving from recently-inserted NUMTs are falsely aligned to mtDNA. With currently used alignment tools, NUMTS that have been in the nuclear genome for a long time are identified as they have accumulated mutations that distinguish them from authentic mtDNA sequences. Only NUMTs that have inserted into the nuclear genome rather recently have not had enough time for distinguishing mutations to occur. As we are therefore not able to directly identify reads arising from recent NUMTs, we instead estimate the probability of incorrectly attributing a NUMT read to the mtDNA genome, using the length distribution of a previously published list of identified NUMTs [28]. For each read insert length, all NUMTs longer than that length were identified as NUMTS from which reads could be generated that could be fully placed within the NUMT. The chance that a read from a NUMT was falsely aligned to the mtDNA genome, given the observed read distribution, was ≤0.06%, and therefore NUMTS were considered to have a negligible impact on mtCN estimation.

We next evaluated the reproducibility of the estimated mtCNs for the two sequencing methods. To estimate standard deviations (SDs) of the methods, intra-individual differences in the chromosomal coverage were evaluated, and the resulting SD estimates were averaged over each tissue and normalized by the average mtCN for the tissue. The shotgun sequencing method returned normalized SD values for chromosomal coverage of <3.2 %, while the SDs of mtCN estimates from capture-enrichment were between 8.6 and 18.2 % (average: 12.6%) (Table 1), indicating more variation in chromosomal coverage of the capture-enriched data (S1 Fig). As both shotgun and capture-enrichment data were available for four tissues (BL, SM, LIV, and MM), we further investigated the reliability of mtCN estimates from capture-enrichment by testing if mtCN estimates from both methods were correlated. While there was a greater enrichment in mtCN for BL than for the other tissues, for all four tissues there was a significant correlation between mtCN values estimated via shotgun sequencing vs. capture-enriched sequencing (Fig 1A). Hence, mtCN estimates based on capture-enrichment sequencing data were considered suitable and used for further investigation into factors influencing mtCNs.

**Fig 1:**
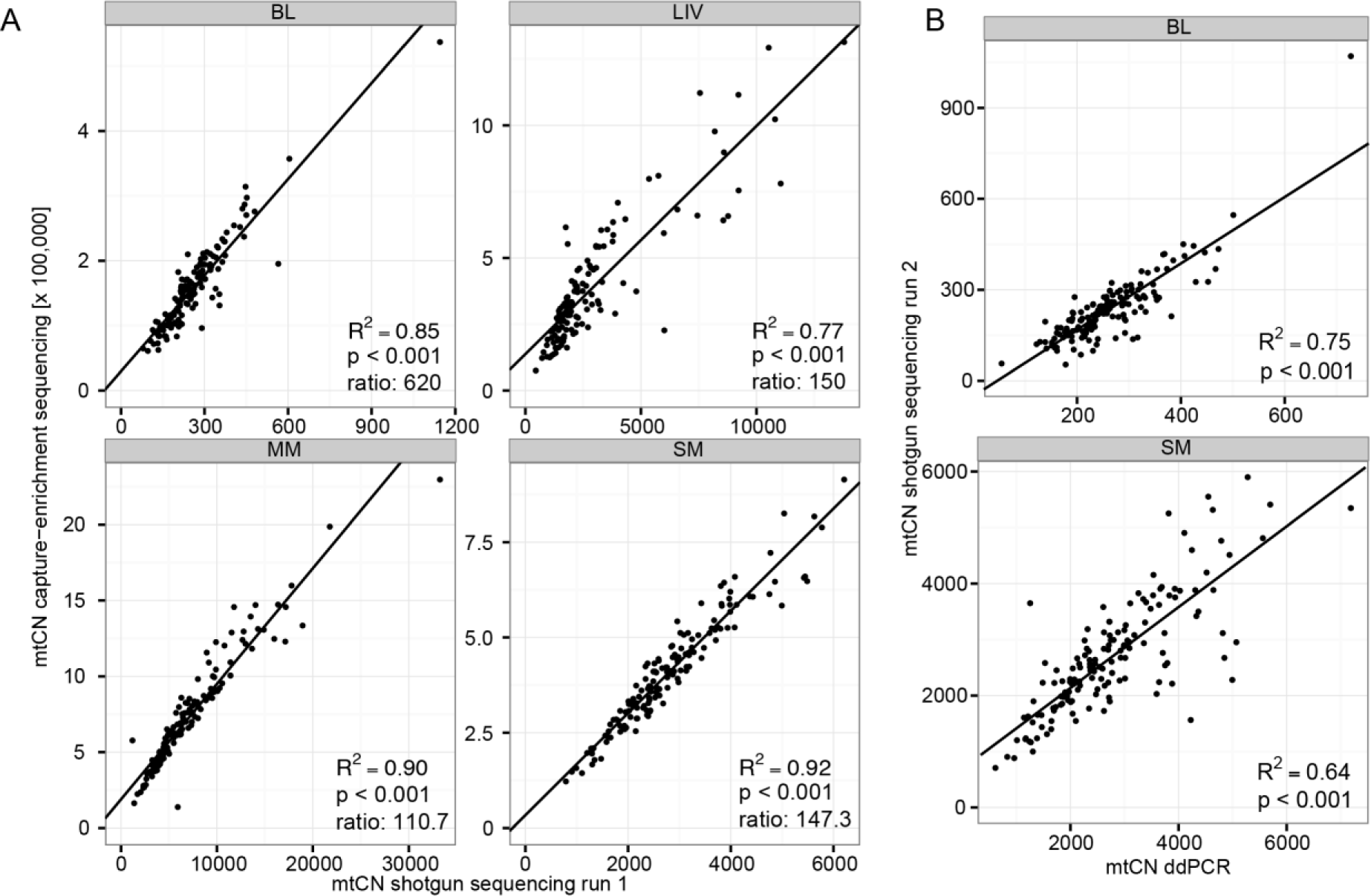
Comparison of mtCN estimates obtained with different methods. Outliers (samples with high SD or samples that resulted in a lack of fit to a normal distribution of the data set) were excluded. R^2^ is the proportion of the variance explained in a linear regression analysis and p is the significance level of the Pearson correlation. (A) MtCN estimates from shotgun sequencing for four tissues (BL, LIV, MM, and SM) vs. those obtained after capture-enrichment. “Ratio” is the amount of enrichment after capture-enrichment, calculated as the average mtCN obtained from capture-enrichment sequencing divided by the average mtCN obtained from shotgun sequencing. (B) MtCN estimates from ddPCR for BL and SM vs. those obtained by shotgun sequencing.

**Table 1:**
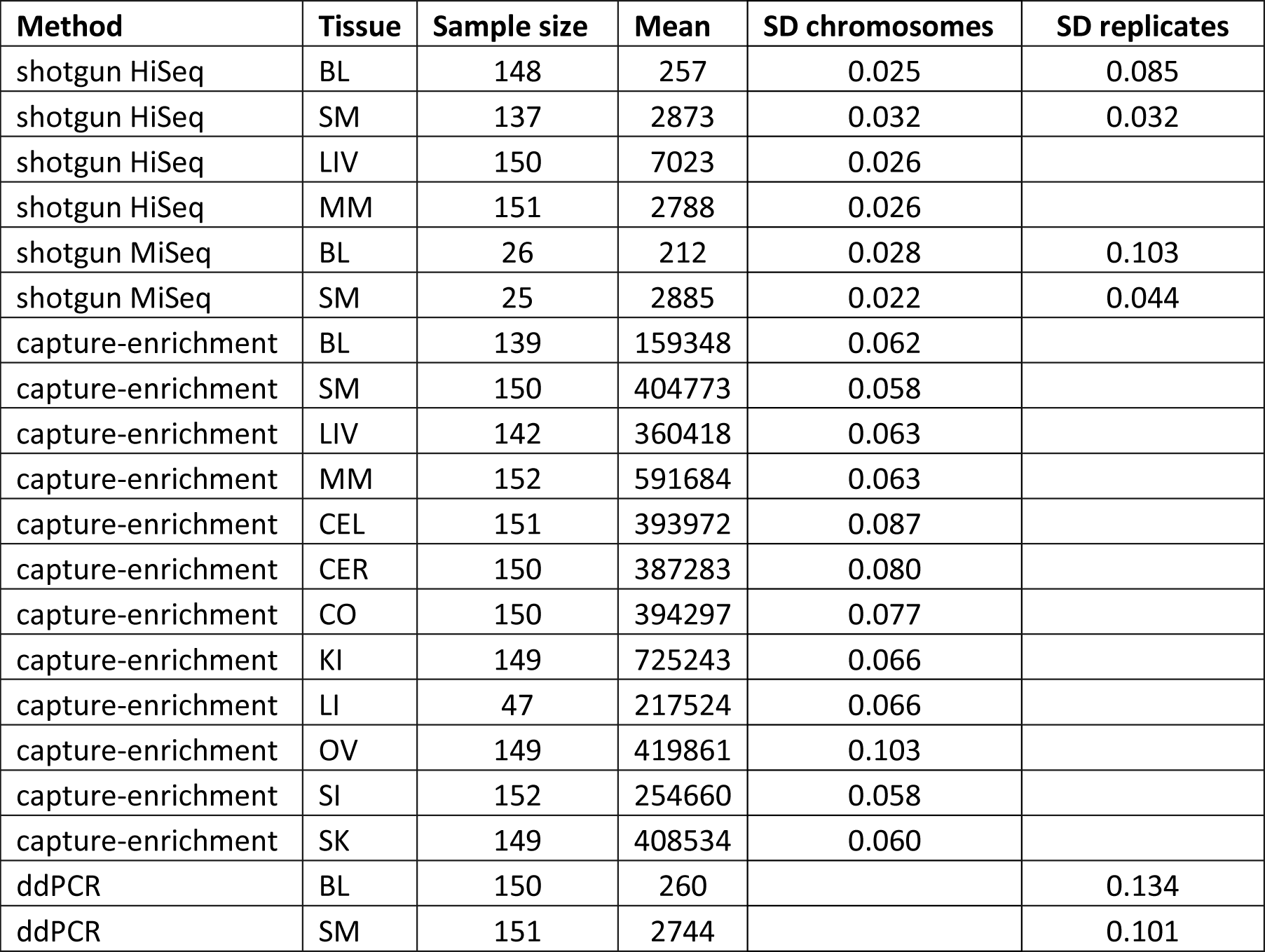
Variation in mtCN estimates from different methods. For each combination of method and tissue, the sample size and mean mtCN estimate across individuals are given. For the sequence-based methods, the mean SD across chromosomes is given (SD-chromosomes); for the shotgun (re-sequencing of the same library and after preparation of new libraries) and ddPCR data from BL and SM tissue the mean SD across replicates for all individuals is given (SD-replicates). All SD values were normalized by the mean mtCN for each tissue.

In a third approach, mtCN was estimated by ddPCR in SM and BL, with each sample measured at least 3 times and the SD calculated over all replicates per sample. Using ddPCR the normalized SD was 10-13% for replicates (Table 1). The estimated mtCN values from ddPCR vs. shotgun sequencing of the same samples were significantly correlated (BL samples: Pearson correlation p<0.001, R^2^=0.75; SM samples: p= p<0.001, R^2^=0.64, Fig 1B) and the mean mtCN values were quite similar (BL: mean mtCN from shotgun sequencing = 257 and from ddPCR = 260; SM: mean mtCN from shotgun sequencing = 2788 and from ddPCR = 2744; Table 1).

To further evaluate the error rates in ddPCR vs. shotgun sequencing, shotgun sequencing was repeated with the same libraries for samples from BL and SM and mtCN and the average SD of the replicates was estimated. The regression coefficient R^2^ was 0.92 for BL and 0.99 for SM (S2 Fig) and the average SD was ≤8.5% and therefore lower than the SD for ddPCR. However, the shotgun sequencing replication was carried out on the same libraries, and hence did not include variation from library preparation or other experimental procedures. To account for this, new libraries were prepared from BL and SM from 26 individuals, sequenced on the MiSeq platform, and the mtCNs were compared (S2 Fig). For both tissues mtCNs could be determined with high reproducibility from shotgun sequencing of an independent library, with R^2^ values of 0.88 and 0.98 for BL and SM samples respectively. As with the ddPCR experiments, the SD of the replicates for mtCN in SM was lower than that for BL (0.044 vs. 0.103 for SM and BL, respectively), indicating a higher reproducibility for the tissue with higher mtCN (Table 1).

### Samples with extremely high mtCN

Remarkably, two samples with extremely high mtCN were identified. One SM sample from a 71 year old male who died of cardiac arrest showed a 137-fold higher mtCN (shotgun sequencing) than the average of all other SM samples (S3 Fig) and a more than 60-fold higher mtCN than the sample with the 2^nd^ highest mtCN. Results from capture-enrichment (32-fold higher than the average, 14-fold higher than the second highest mtCN) and ddPCR (80-fold higher than average, 30-fold higher mtCN than the second highest) were similar. In addition, one LIV sample from a 72 year old male who died of multiple organ failure showed a 20-fold higher mtCN than the tissue average in shotgun sequencing (8-fold in capture-enrichment). The mtCN estimates for other tissues in these two individuals are all within the average range for that tissue, indicating that this extreme increase in mtCN is tissue-specific.

In other tissues, the highest mtCN value was maximum 4-fold higher than the average in shotgun sequencing (6-fold in capture enrichment), showing that extremely high mtCNs are rare in the data set.

### Inter-individual differences in mtCN and correlations of mtCN with age, sex and haplogroup

For each method, we tested if mtCN estimates were correlated between pairs of tissues within an individual (Fig 2). All p-values were corrected for multiple testing using Benjamini-Hochberg correction [31]. In the capture-enriched data, positive correlations were identified between SI and LI (r=0.31, p<0.001), between CO and CER (r=0.33, p<0.001) and between KI and CER, CO, SI and SM. In addition, mtCN in SK was negatively correlated with that in SI, LI and in LIV. In the shotgun sequencing data there was a negative correlation of mtCN in SM vs. LIV (Pearson’s r=-0.23, p=0.042, Fig 2), but this was not observed in the capture-enriched data (Fig 2). Overall, we did not observe any regular pattern in the intra-individual variation in mtCN among different tissues (S4 Fig).

**Fig 2:**
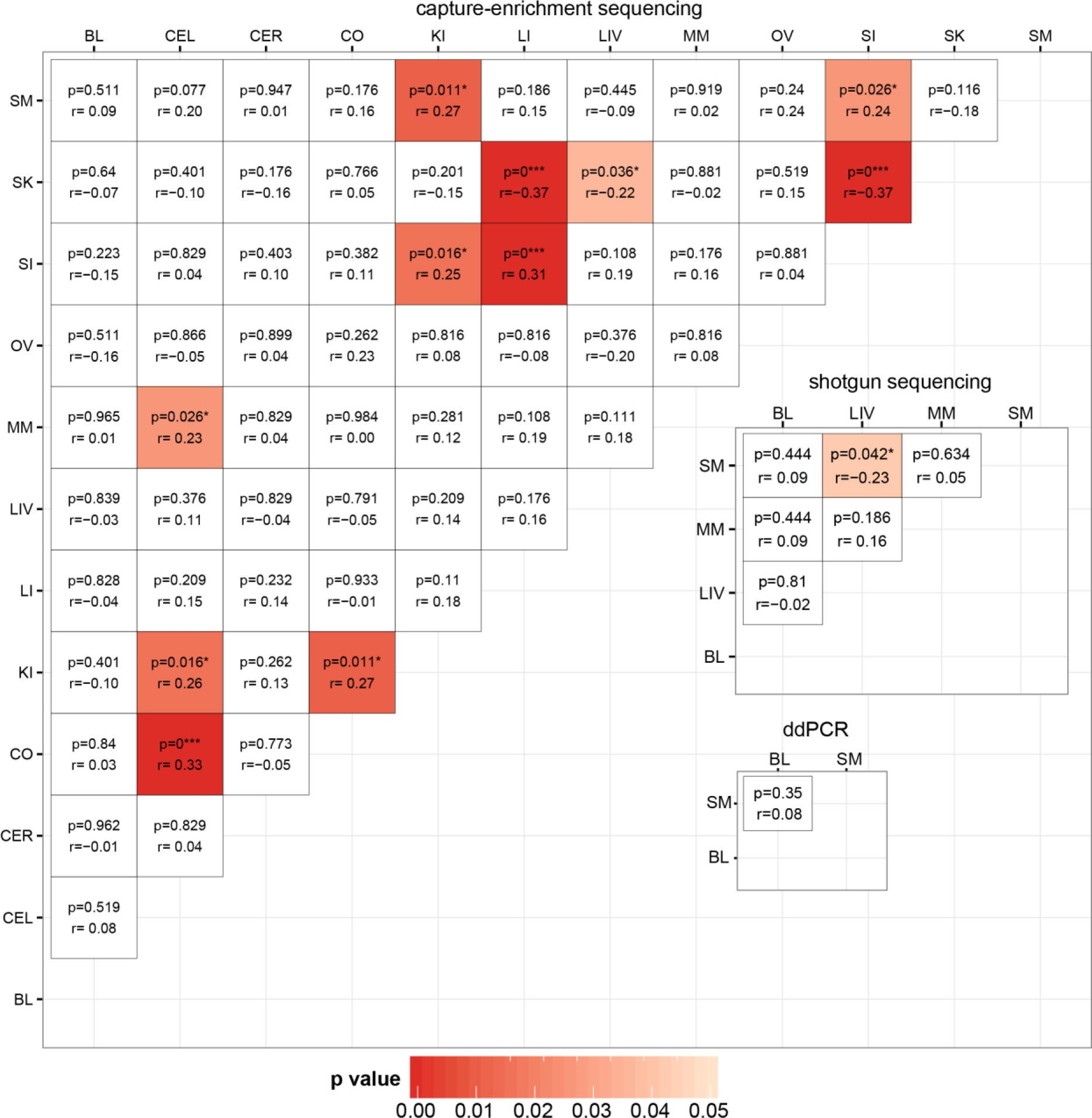
Correlation analysis of mtCN estimates from different tissues by three different methods. Corrected level of significance (p) and the Pearson correlation coefficient (r) are given in each field; fields are colored by p-value according to the scale.

We also investigated associations between mtCN and age, sex, or haplogroup. There was a strong age-related decrease of mtCN in SM (r=-0.25, p=0.016 in shotgun sequencing, Fig 3A, S2 Table). Remarkably, this association holds only for males (r=-0.35, p=0.008 in males compared to r=-0.15, p=0.442 in females; S2 Table). Similar correlation coefficients were obtained for mtCN estimates from capture sequencing and ddPCR, although the p-values were not significant after correction for multiple testing (S2 Table). In LIV, mtCN shows an increase with age that is significant in capture enrichment data (r=0.27, p=0.022) and approaches significance in shotgun sequence data (r=0.20, p=0.088; S2 Table). This increase was mainly due to a subset of individuals (Fig 3A) with higher mtCNs. When looking at samples with mtCN≥4,500, these showed a stronger correlation of mtCN with age (r=0.64, p=0.016) than the group mtCN≤4,500 (r=0.19, p=0.048).

**Fig 3:**
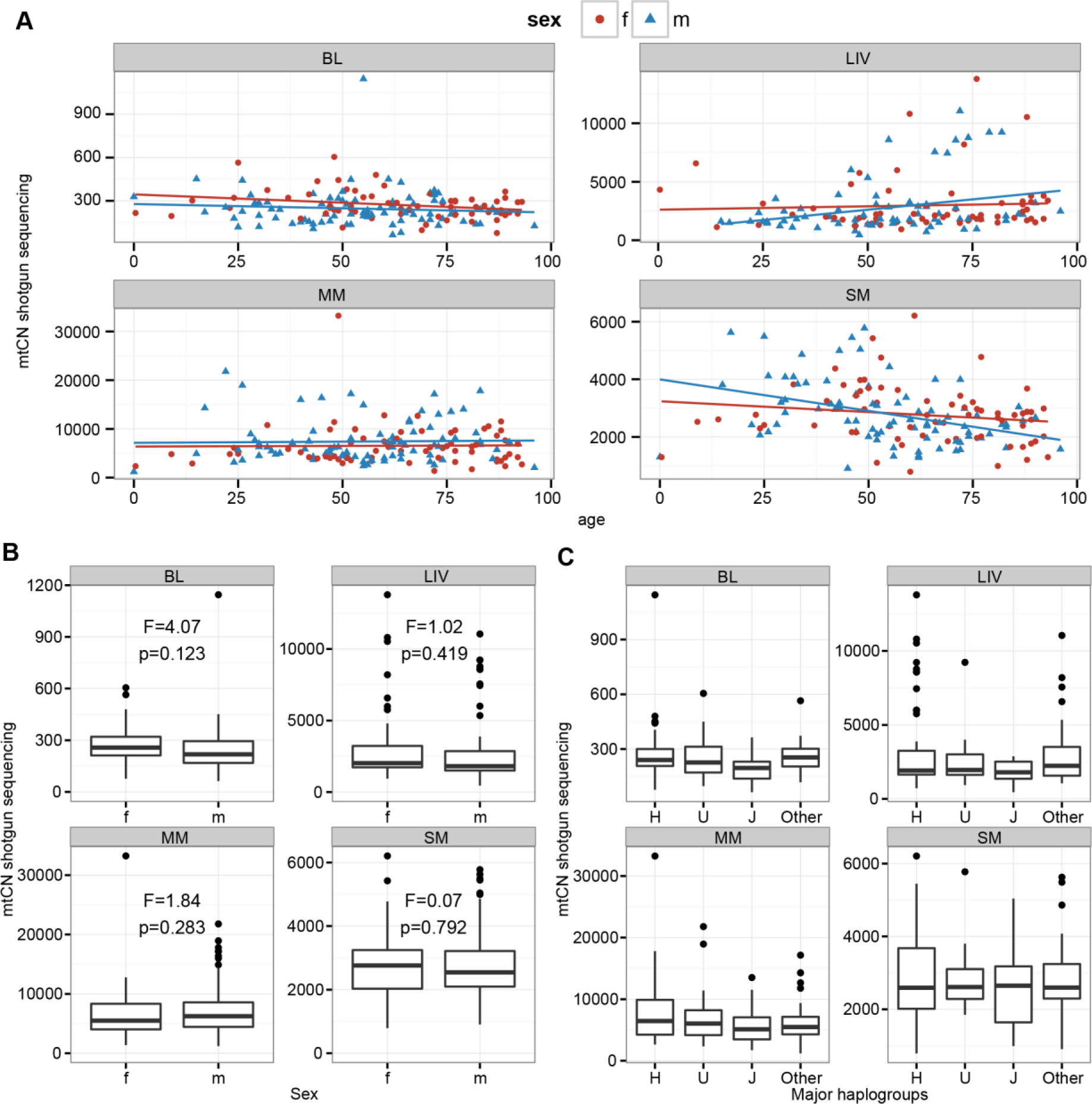
Correlation analysis of mtCN from shotgun sequencing with age, sex and haplogroup. Males (m) and females (f) are indicated. (A) Correlation of mtCN with age. (B) Correlation of mtCN with sex. F- and p-values are shown for each tissue. (C) Correlation analysis of mtCN with haplogroup, identified as H, J, U or other haplogroup.

No solely sex-related associations with mtCN were identified for any tissue (Fig 3B, S2 Table) and there were no significant associations between mtCN and major haplogroup (Fig 3C, S3 Table).

### Associations between mtCN and heteroplasmy

As all samples in this study were previously analyzed for heteroplasmy [19], we tested for associations between mtCN and the total number of heteroplasmic positions as well as the MAF at specific heteroplasmic positions. SM exhibited a highly significant negative correlation between mtCN and the total number of heteroplasmies for all three methods (r=-0.27 to -0.31, all p≤0.002; Fig 4, S4 Table). BL and LIV also exhibited significant negative correlations between the total number of heteroplasmies and mtCN, but not for all of the methods (S4 Table). As the number of heteroplasmic sites has been shown to significantly increase with age [16, 19], we analyzed the influence of age and the total number of heteroplasmic sites on mtCN in SM by partial regression. MtCN was best explained by age (p<0.05), although the number of heteroplasmies also showed a slight correlation with mtCN that was borderline significant for shotgun sequencing data (p=0.049, Fig 4). In LIV, changes in mtCN were significantly associated with age (p<0.05, Fig 4), whereas BL showed a higher partial regression of mtCN with heteroplasmy than with age (Fig 4). Overall, these results indicate that correlations of mtCN with the total number of heteroplasmies are explained mainly by age rather than the actual increase in the number of heteroplasmic sites.

**Fig 4:**
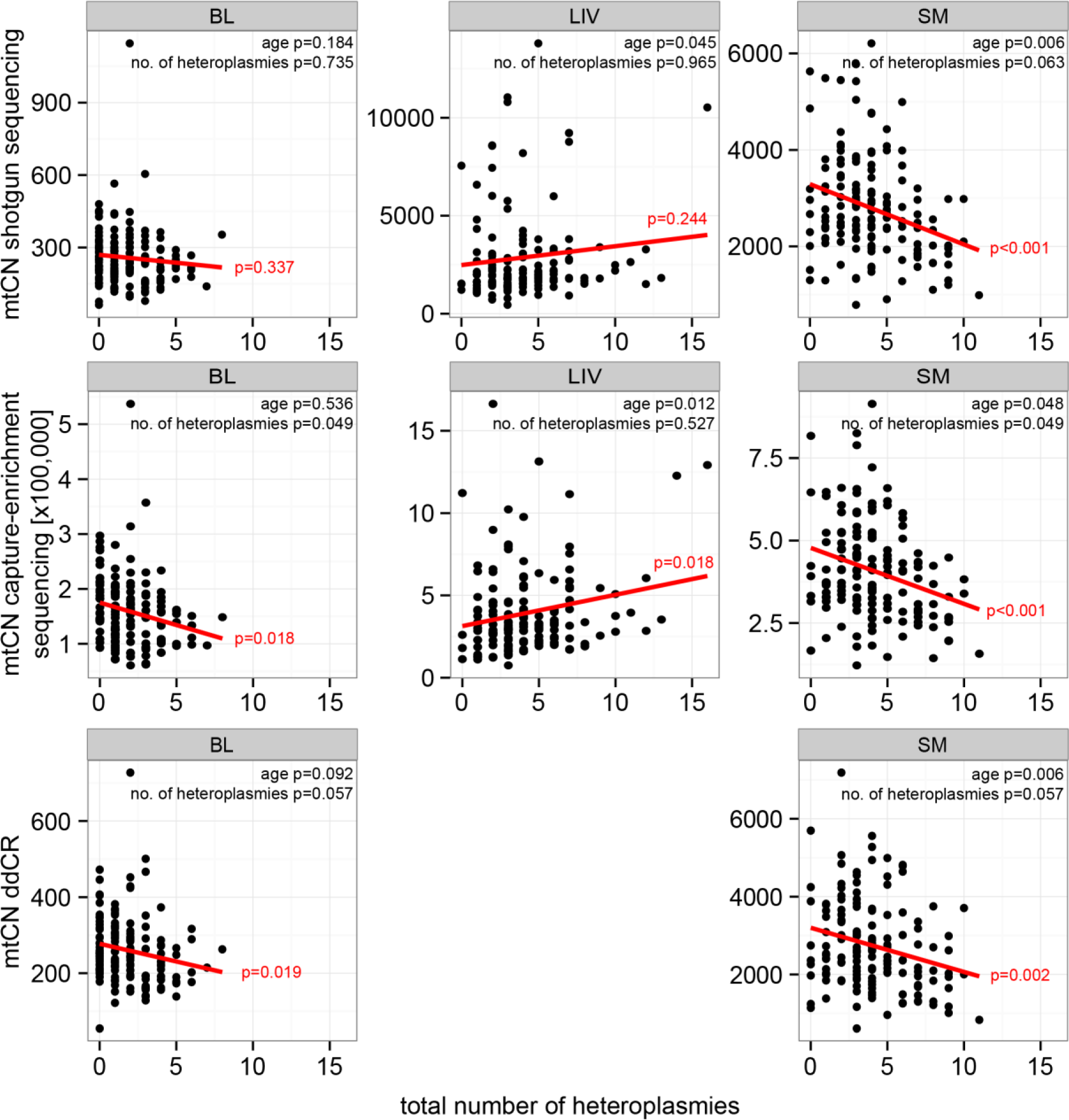
Correlation analysis of mtCN from shotgun sequencing with the number of heteroplasmic sites in BL, LIV and SM for shotgun sequencing, capture-enrichment and ddPCR. P-values of linear regression and regression line are given in red. P-values for partial regression of mtCN with age and the total number of heteroplasmies are shown in black.

With respect to associations between mtCN and the MAF at single heteroplasmic sites, analyses were limited to positions that were heteroplasmic in at least ten individuals for a tissue (S5 Table). After correction for multiple testing, sites 408 and 16327 each showed a significant association between mtCN and MAF in SM for shotgun sequence data (p=0.014 for both sites, Fig 5, S6 Table). While the consensus allele at position 408 is a T in all tissues and individuals in this study, an A allele is observed as a heteroplasmy (MAF = 2-28.2 %) in 65/152 individuals in SM. The consensus allele at position 16327 is a C in all but one individual in this study, but heteroplasmy involving a T (MAF = 2-20%) occurs in SM in 46/152 individuals. Other alleles were not found at either position in these samples [19]. As the MAF at these positions also increases with age [19], we applied partial regression to test whether age or heteroplasmy was more strongly associated with mtCN. While age was again significantly correlated with mtCN (p=0.001 for both sites), the MAF at both heteroplasmic sites also showed significant associations with mtCN (p=0.022 (408), p=0.02 (16327), Fig 5). The total number of heteroplasmies, however was not significantly associated with mtCN (p>0.05). This indicates that changes in mtCN in SM are associated with age, with the MAF at positions 16327 and 408 also explaining some of the change in mtCN.

As the aging-dependent decline in mtCN was significant in males but not in females, we asked whether there were sex-dependent differences in the correlation between mtCN and MAF, too. After correction for multiple testing, neither MAF at site 408 nor at site 16327 was significantly correlated with mtCN in males or females alone (males: 408: p=0.64, 16327: p=0.64; females: 408: p=0.4, 16327: p=0.4).

**Fig 5:**
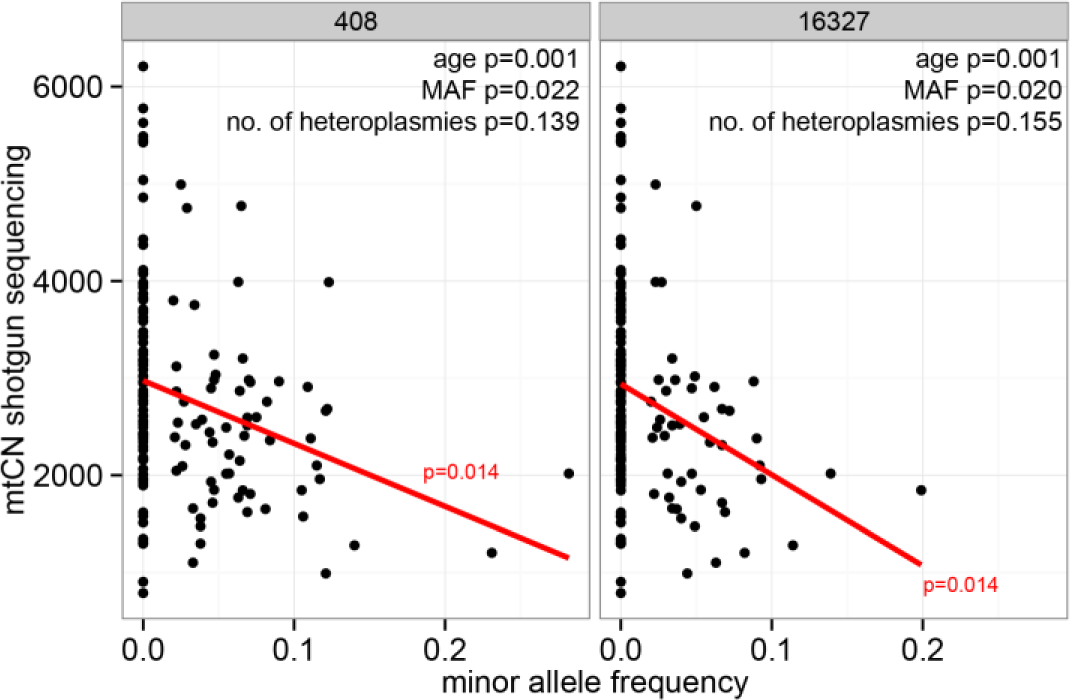
Correlation of mtCN with MAF at positions 408 and 16327 in SM. P-values of linear regression and regression line are given in red. P-values for partial regression of mtCN with age, MAF and the total number of heteroplasmies are shown in black.

No stronger correlations were found when testing the MAF of pairs or tripletons of heteroplasmic sites, indicating that heteroplasmy at two or three specific sites was not more associated with variation in mtCN than heteroplasmy at single sites. In sum, mtCN in SM differed dramatically from other tissues in exhibiting highly significant age, sex and heteroplasmy-dependent associations.

## Discussion

### Methods for mtCN determination

In this study, mtCN from different tissues was estimated using three different methods. Exact measurements of mtCN can be complicated in some tissues as intra-individual differences in mtCN of a single tissue can occur. These differences arise from mosaic-like structures as those found especially in myocardial muscle [4] and one should therefore focus investigations of mtCN on rather homogenous tissues.

In ddPCR and shotgun sequencing experiments, SM samples showed smaller SDs for multiple measurements than BL samples. This could indicate that the precision of the two methods is increasing with higher mtCNs as small absolute measurement errors have less impact on the final results. In general, the shotgun sequencing method returned normalized SD values that were smaller than those for ddPCR. The high variance between measurements in ddPCR might arise from the several pipetting steps for mtDNA- or nDNA-specific primers as well as additional dilution steps between nDNA and mtDNA-determination. In Illumina shotgun sequencing, the only step in which mtDNA and nDNA compete is the loading of DNA molecules into the nanowells of the flowcell. The distribution of reads across chromosomes indicates that for most chromosomes this happens randomly without any bias for certain sequences. Chromosomes that had to be excluded from the data set showed a large SD across all samples due to the assembly of reads with lower mapping quality. Since shotgun sequencing exhibited high reproducibility of mtCN with two independently prepared libraries and moreover allows high-throughput analysis of large sets of samples, this is the preferred method for mtCN determination.

We assumed that capture-enrichment linearly increases the amount of mtDNA without saturation effects when estimating mtCN from sequencing after capture-enrichment. However, we observed that the enrichment process is non-linear, resulting in a stronger enrichment of mtDNA in samples with smaller mtCN. Samples with very high mtCN, like the SM-sample with a 100-fold increase in mtCN, already have a very high proportion of mtDNA prior to enrichment and will therefore be underestimated after enrichment. Samples with lower mtCN, like BL samples, have a low starting ratio that is strongly increased in the enrichment process leading to overestimated mtCNs after enrichment. Hence, enrichment most strongly influences the tails of the mtCN distribution; however, the overall high correlations between mtCN estimates from capture-enriched vs. shotgun data indicates that mtCN estimates by the former method do reliably reflect relative mtCN values.

### Occurrence of samples with very high mtCN

We identified two samples from different individuals that had 20-fold (LIV) and a ≈100-fold (SM) higher mtCN compared to the tissue average. Similar results were obtained with the different methods for mtCN determination, indicating that these high mtCN are not method-dependent errors. To our knowledge no such increases in mtCN have been described before. A strong increase of mtCN has been described when large parts of the coding region of the mtDNA are deleted [33] and steep increases in copy number occur during cell differentiation, such as an 1100-fold increase in mtCN between day 5 and 6 of differentiation of pluripotent embryonic stem cells [34]. In addition, external factors such as stress [35] or diseases such as cancer, diabetes or HIV (reviewed in [36]) can increase mtCN. Cai *et al.* reported a stress-induced increase in mtCN in liver, but a decrease in SM in mice. Interestingly, mtCN in those two tissues was negatively correlated in our shotgun sequencing data. As we do not have any information on the stress levels of the individuals in our sample set, no further comparison of stress levels with mtCN is possible. However, previous studies found that stress increased mtCN by only around 2-fold [35], which cannot explain the samples with very high mtCN found here.

An excess of replication has been associated with several mtDNA regulatory proteins, such as TFAM [6] or PGC-1, a common transcriptional co-activator of nuclear receptors [37]. The individuals we identified did not suffer from any diagnosed disease that might impact mtCN, and these extreme mtCNs occurred in just one tissue in each individual. Heteroplasmy, haplogroup or age do not explain this extreme increase in mtCN. As all enzymes required for mtDNA replication control, like TFAM or PGC-1, are encoded in the nucleus [24], we speculate that mutations in nuclear-encoded genes might trigger this increase; however an environmental cause cannot be excluded.

### Regulation of mtCN in tissues

Following mtCN estimation, several parameters were tested for possible correlations with mtCN using linear regression and a Pearson correlation test. The data from capture-enrichment returned the fewest significant results, which probably arises from Benjamini-Hochberg corrections for multiple testing as for capture-enrichment there were 11 or 12 tissues (ovarian tissue data could only be analyzed in females) analyzed compared to four in shotgun sequencing and two in the ddPCR. As the F- and r-values, which were not corrected for multiple tests, were in the same range for all three methods, we conclude that the most striking correlations were identified regardless of the method used.

To our knowledge, few studies have investigated differences in relative mtCN between different tissues of an individual; however, one previous study did investigate mtCN in samples from brain, SM and MM from 50 individuals [11]. This study identified correlations of mtCN between most tissues. In our study, different tissues of the same individual showed a high variation in mtCN, not only for the total number of copies, but also for the relative number with respect to the tissue average. Positive correlations were mainly found for similar tissues, such as different parts of the brain or for large and small intestine. Interestingly, mtCN in skin had a strong negative correlation with mtCN in both intestinal tissues (as well as liver). Skin and intestine are surface tissues with exposure to different microbiomes [38], and hence different environmental circumstances might be influencing mtCN. The variation in mtCN between different tissues of an individual is largely uncorrelated (Fig 2). Thus, it is not the case that individuals tend to have generally high or low mtCN values across all tissues, but rather mtCN varies in a tissue-specific fashion. This is in good agreement with a recent study investigating stress-induced changes in mtCN [35], which showed that stress increased mtCN in liver, but decreased mtCN in muscle in mice. Therefore, external factors can produce tissue-specific changes in mtCN.

Specific haplogroups have been suggested to influence mtCN. For example, a comparison of cybrid cells from haplogroups H and Uk found that cells of haplogroup Uk exhibited lower mtCN than those of haplogroup H [39]. In our present study, no individual belonged to haplogroup Uk. Another study comparing haplogroups H and J found that the haplogroup J defining mutation increased TFAM binding and as a consequence mtDNA replication and mtCN [40]. In our data set no significant differences in mtCN were detected for different haplogroups, including haplogroup J. However, the relatively small sample size for each haplogroup in our study (e.g., only 14/152 individuals with haplogroup J) means that we lack power to identify potential haplogroup-related effects on mtCN, and further studies with larger sample sizes are warranted.

Age is another factor that could influence mtCN; for example, previous studies have described age-dependent decreases in mtCN for BL in individuals older than 50 years [8, 10]. Although in our study all three methods exhibited a negative correlation between age and mtCN in BL for individuals over 50, only for ddPCR did the correlation approach significance (p=0.09, S2 Table). In contrast, the age-dependent decrease in SM is very striking in our data, especially in males. This result seems to contradict other studies of mtCN in the same tissue [4, 41]. However, the study of Barthelemy *et al.* investigated samples from younger individuals (2-45 years, n=16), and no information on the sex distribution of the individuals was given in either study. The strong decrease in mtCN with age in males may be explained by the composition of muscle tissues. Males tend to have a higher ratio of fast-contracting muscle fibers (type IIB) over slow fibers (type I) [42]. Slow muscle fibers are characterized by a high activity, strong coupling of the electron chain and therefore high oxygen capacity. To account for aging effects, slow fibers reduce the coupling of the electron chain, resulting in low reactive oxygen species and stable mitochondrial function in old age [43]. Fast contracting muscle fibers, on the other hand, have a short longevity and are susceptible to aging. They show an advancing transformation to slow-contracting muscle fibers during aging which is accompanied by a reduction in mtDNA content [42]. Due to these differences in muscle composition, effects on mtCN during aging might be stronger in males than in females.

Finally, we investigated the influence of heteroplasmy on mtCN. The total number of heteroplasmic sites was strongly negatively correlated with mtCN in SM in our data. As the number of heteroplasmic sites is also strongly age-related [16, 19] and mtCN in SM is decreasing with age, the major predictor of mtCN remains unclear after linear regression analysis. We tried to account for this by partial regression, in which one factor is set constant while the effect of the other is investigated. For the total number of heteroplasmies, we found that age had a more significant correlation with mtCN, indicating that a non-specific enrichment of heteroplasmic sites is not the major effector for changes in mtCN.

We then investigated potential associations between the MAF at specific heteroplasmic positions and mtCN. Previous studies have found that sequence variations in the polycytosine tract at positions 16180-16195, and in a TFAM binding motif at position 295 in the mtDNA control region, are associated with changes in mtCN [5, 40]. Another study on saliva from women that suffered from major depressive disorder showed that heteroplasmy at site 513 was significantly correlated with changes of mtCN [44]. In our study, we also found indications for associations between the MAF at common low-frequency heteroplasmic sites (408 and 16327) and mtCN in SM. The MAF at both positions has been shown to be highly correlated with age [18-21, 45]. Heteroplasmy at position 16327 is highly specific for SM as it was not found in more than four individuals in any other tissue in our previous study [19]. Heteroplasmy at position 408 is even more common in SM, but was also found in some individuals in CER (12/142) and CO (13/152, S5 Table) with very low MAF (≤1.1%) in the previous study [19]. Together with site 189, site 408 was reported to be associated with aging-dependent decrease mtCN in 52 females from family units [46]. However, the authors of this study stated that the modest changes in MAF that they observed at those sites might not be the major determinant of mitochondrial dysfunctions during aging [46]. In our data set, we found that the correlation of MAF at site 408 with mtCN in either males or females alone was not significant after correction for age. The reduced sample size (84 males and 67 females instead of 151 in the entire sample set) might reduce the power to detect significant effects in the data sets.

Site 408 is adjacent to the transcription start site within the light strand promoter (S7 Fig), which initiates production of an RNA primer for heavy strand replication [24], while 16327 is located within the TAS-region (S7 Fig), which regulates D-loop formation. The ratio of D-loops over double stranded DNA has been shown to be crucial for mtCN control [47] and parts of the TAS region, including site 16327, have been identified as a putative binding site for unknown proteins [48]. As the occurrence of heteroplasmy arises in an allele-specific way, we hypothesize that heteroplasmy at positions 408 or 16327 might lead to changes in replication regulation. The fact that MAF at 408 was not correlated with mtCN in CER and CO might be explained by the much lower MAF in these tissues.

Intra-individual deleterious mutations in mtDNA may persist at a low level due to a lack of negative selection [22, 49] as they need to reach a critical frequency threshold in order to impact mitochondrial function [50]. If the negative selection pressure is relaxed in one tissue relative to others, due to the metabolic needs of that tissue (especially during aging) then tissue-specific deleterious heteroplasmies could potentially increase in frequency due to this relaxation of functional constraints [51-53]. Alternatively, low-frequency mutations might also rise in frequency via positive selection, if they are advantageous for that specific tissue [15, 19], e.g. if they influence replication rates [16, 54]. The results of the present study do not bear on the reason(s) for the tissue-specific and age-related increases in heteroplasmy. However, reduced mitochondrial activity does lead to reduced accumulation of reactive oxygen species and protects mitochondria against aging [55]. We hypothesize that positive selection for a heteroplasmic allele could arise via a slight decrease in mtCN that results in a slowdown of mitochondrial metabolism. This, in turn, would reduce the production of aggressive reactive oxygen species and hence provide a better cellular fitness during aging, in accordance with the “survival of the slowest” hypothesis [56].

Both alternative minor alleles at these two positions (408A and 16327T) are fixed in other haplogroups than those investigated here [57]. For example, 16327T is one of the alleles defining haplogroups C and U1B. One individual in our data set was identified as haplogroup U1B, but did not show any substantial difference in mtCN compared to other individuals (S7 Table). While one individual does not allow relevant interpretation of haplogroup-dependent changes in mtCN, it does suggest that overall the 16327T allele is not highly deleterious. However, functional differences between haplogroups as well the functional impact of reduced mtCN in aging SM remain to be elucidated.

In conclusion, we found that SM exhibits several interesting properties with respect to mtCN and heteroplasmy, including age-related decreases in mtCN in males, and decreases in mtCN associated with increasing MAF at two heteroplasmic sites in the control region. Several additional questions remain. For example, why do the minor alleles at tissue-specific heteroplasmic sites remain at rather low levels and do not reach higher and therefore presumably more detrimental frequencies in the cell during aging? One effect of aging is the reduction of mitochondrial fusion and fission events that allow an exchange of genetic material between mitochondria in young individuals [58]. It has been proposed that reduced fusion and fission prevent cells from spreading detrimental mitochondria throughout the cell [59] and therefore decelerate aging effects. This might also explain the persistence of low frequency of heteroplasmy in aging individuals.

Several sites in the control region show site-specific and allele-specific heteroplasmy in specific tissues, such as position 60 and 94 in LIV and KI or position 204 and 564 in MM [19], yet in this study only the heteroplasmy at sites 408 and 16327 in SM showed a correlation with mtCN. Overall, elucidating the functional consequences of tissue-specific heteroplasmy remains a major challenge for further investigation.

## Acknowledgments

We thank Janet Kelso, Matthias Meyer and Martin Petr for advice and supportive discussions on mtCN estimation from sequencing data; Michael Dannemann for help with statistics and comments on the manuscript; and Roland Schröder for technical support.

## Supporting Information

**S1 Fig:**
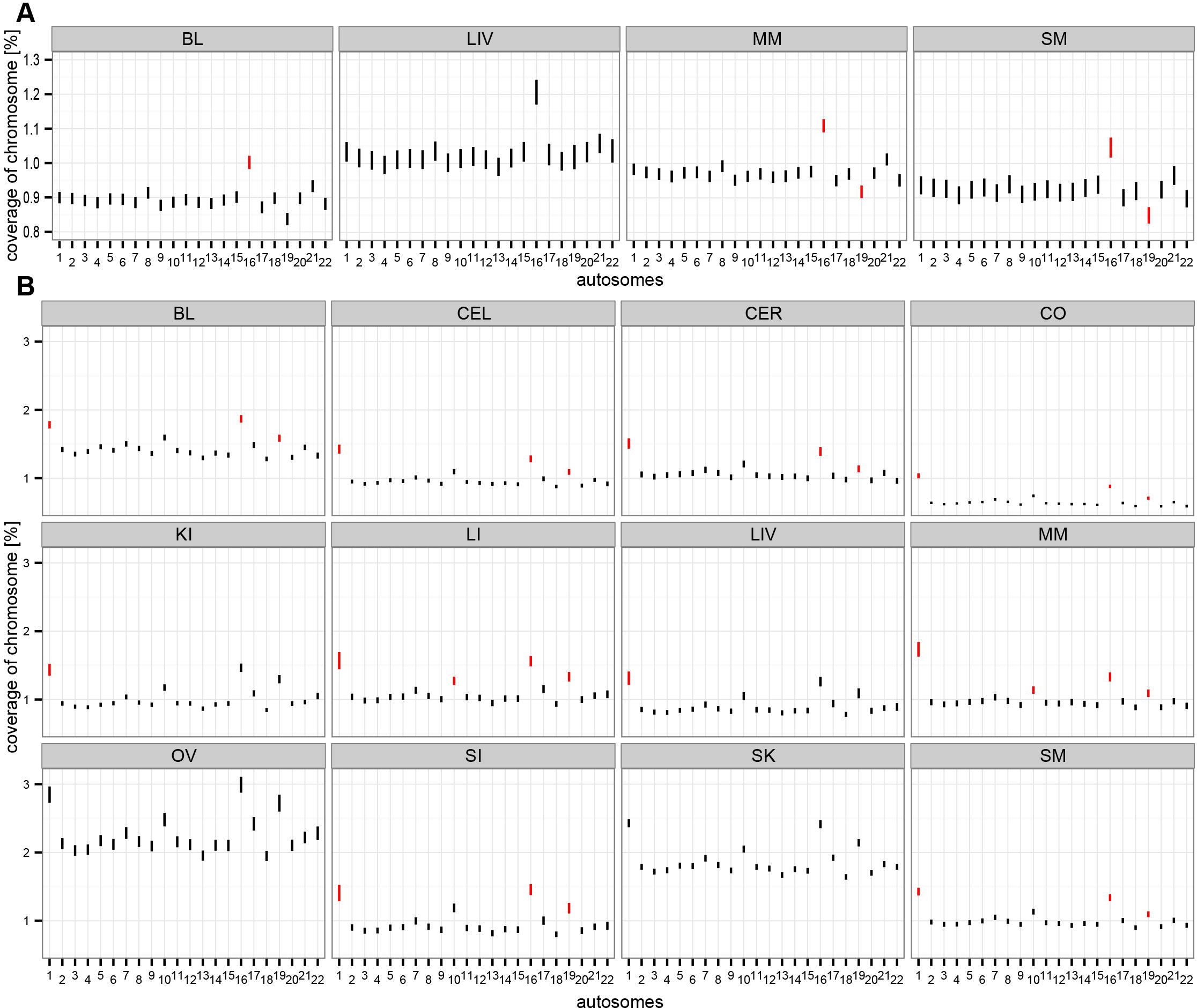
Average autosomal coverage and SD of coverage per tissue per chromosome (A) in shotgun sequencing data (B) in capture-enrichment sequencing data. Each bar indicates the 95% confidence interval of the chromosomal coverage. Red bars indicate significant outliers according to SD.

**S2 Fig.**
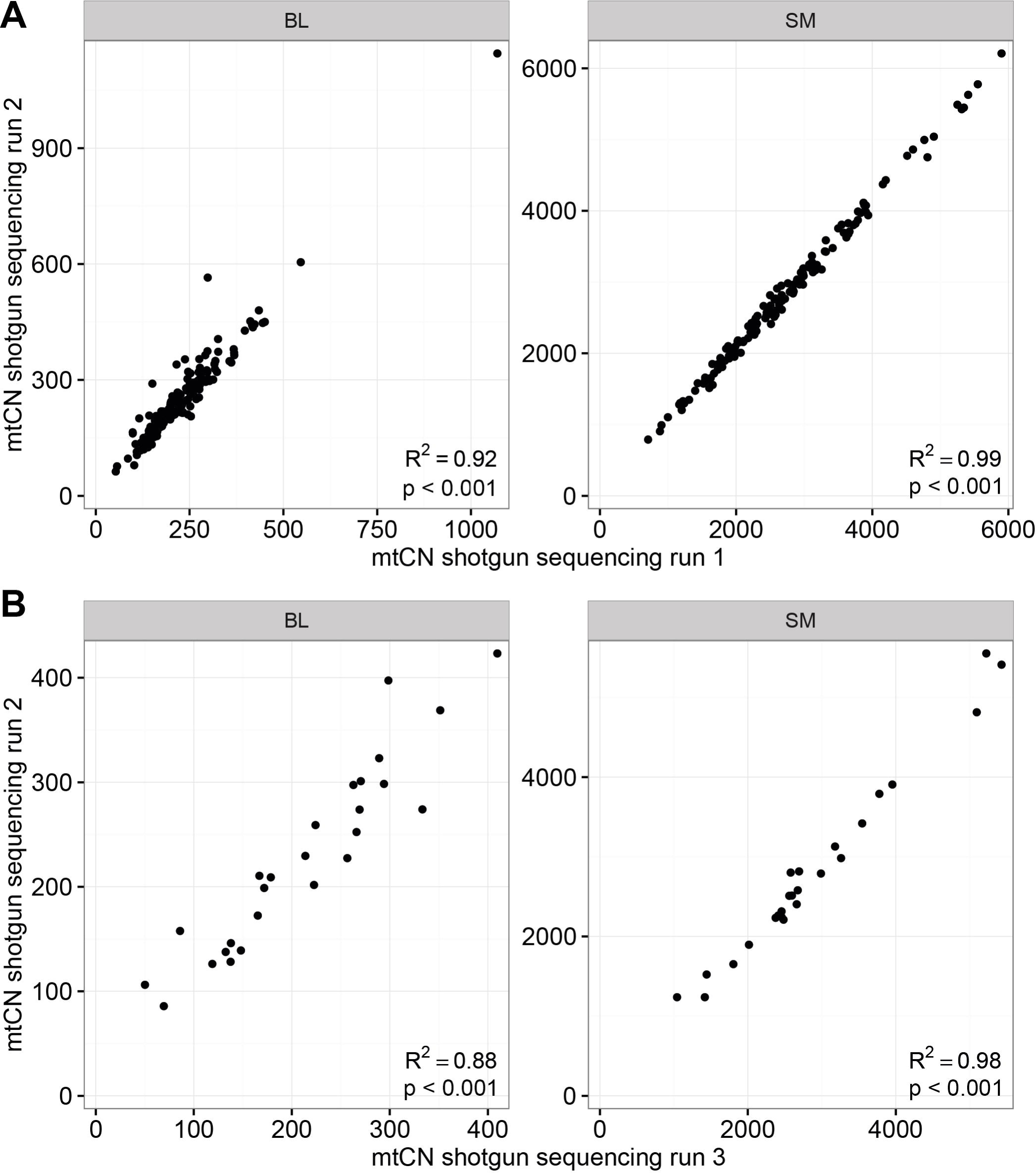
Reproducibility of mtCN determination using shotgun sequencing data. **A** BL and SM samples were sequenced in two independent sequencing runs of the same libraries. **B** independent libraries were prepared from a subset of 26 individuals for BL and SM samples and sequenced on a MiSeq, mtCNs from this sequencing run are plotted against the corresponding mtCNs from the first library. R^2^ represents the coefficient of determination of a linear regression analysis adjusted for the degrees of freedom.

**S3 Fig.**
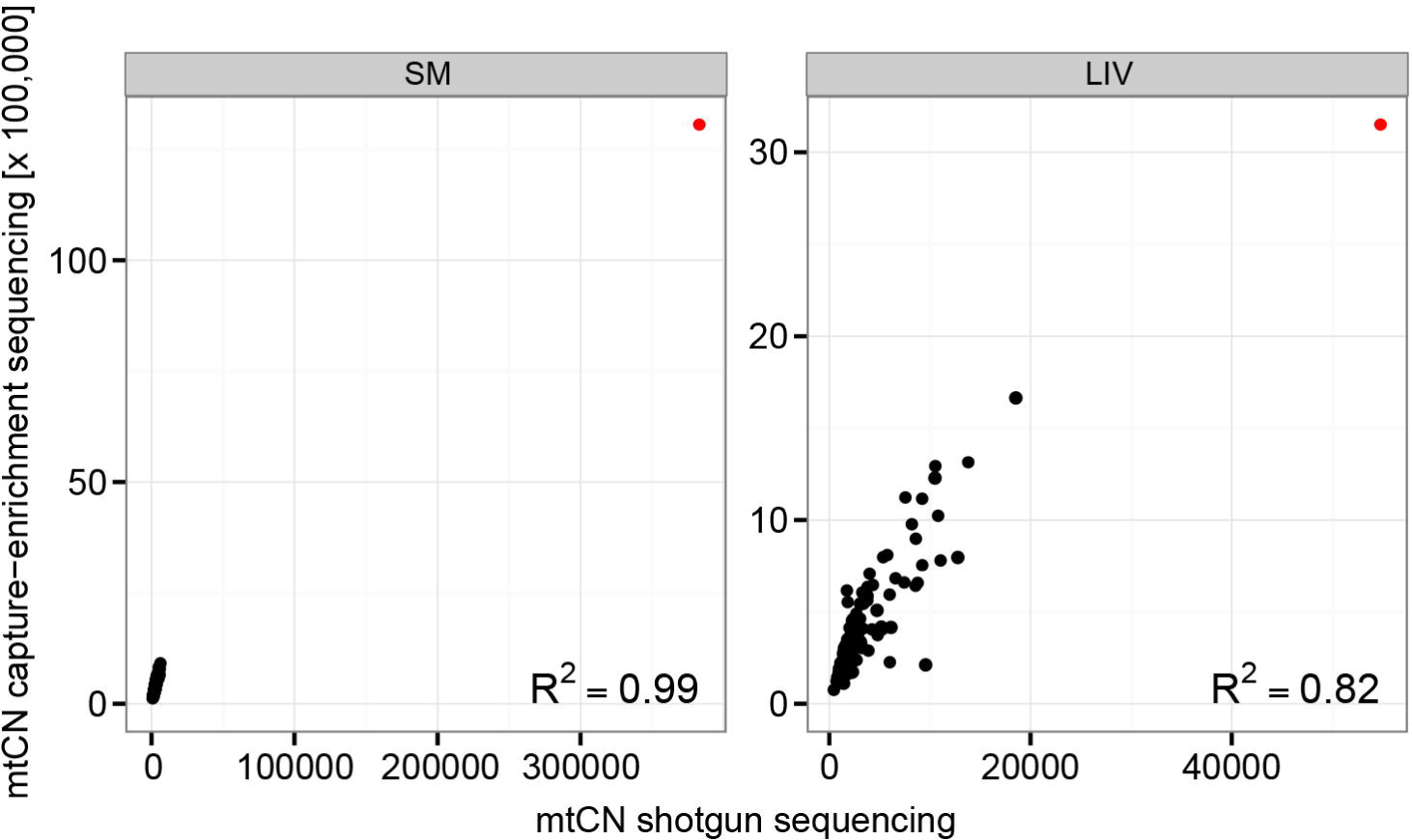
MtCN estimated by shotgun sequencing vs. mtCN estimated after capture-enrichment sequencing, including outliers. One SM sample and one LIV sample (red) had very high mtCNs, and were excluded from the correlation analyses for violating the assumption of a normal distribution.

**S4 Fig.**
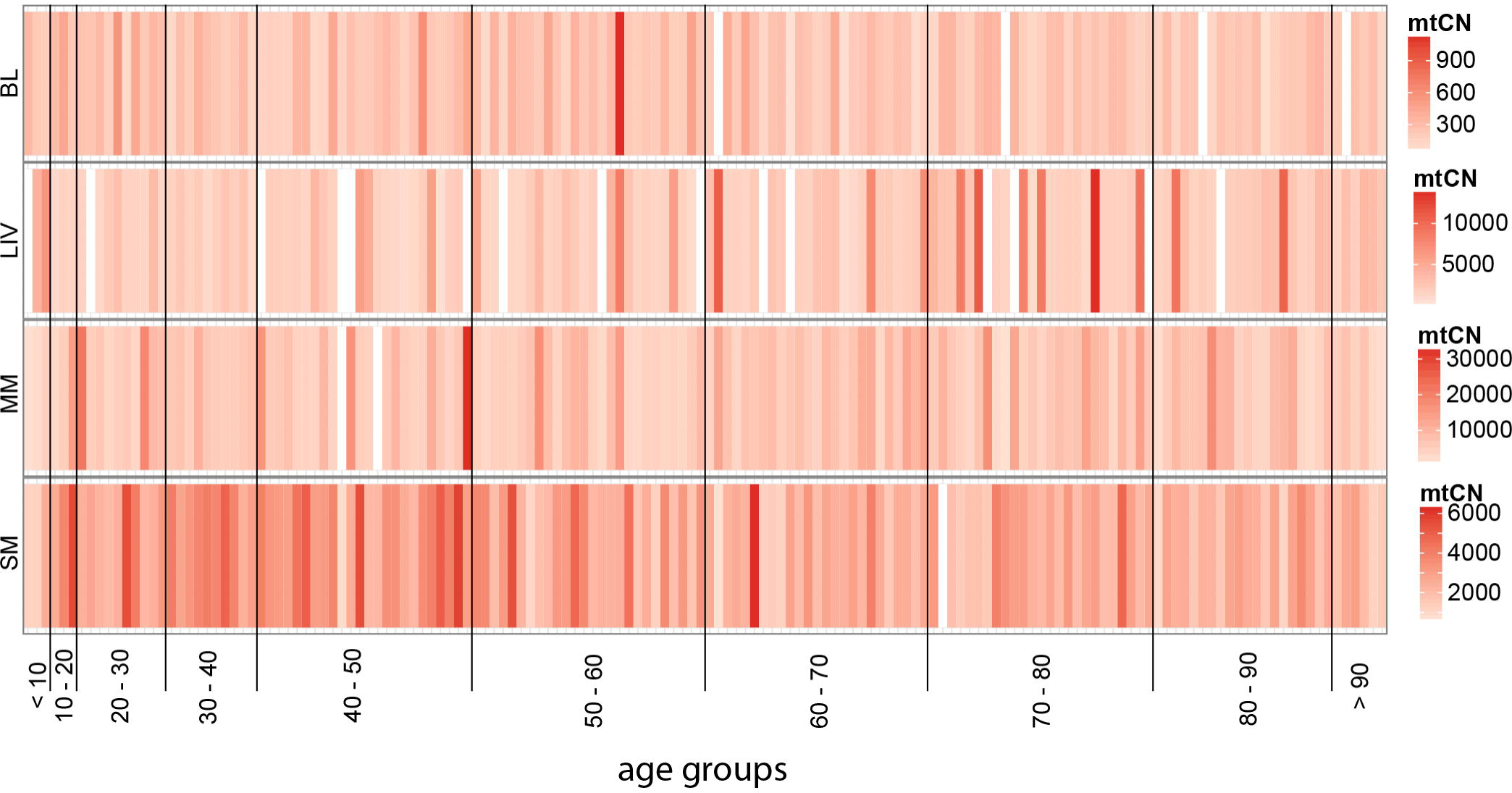
Heat plots of relative mtCN of single individuals in BL, LIV, MM and SM estimated by shotgun sequencing. Vertical bars indicate single individuals, sorted from young (left) to old (right). Coloring of a vertical bar indicates the mtCN according to the scale on the right of each plot.

**S5 Fig.**
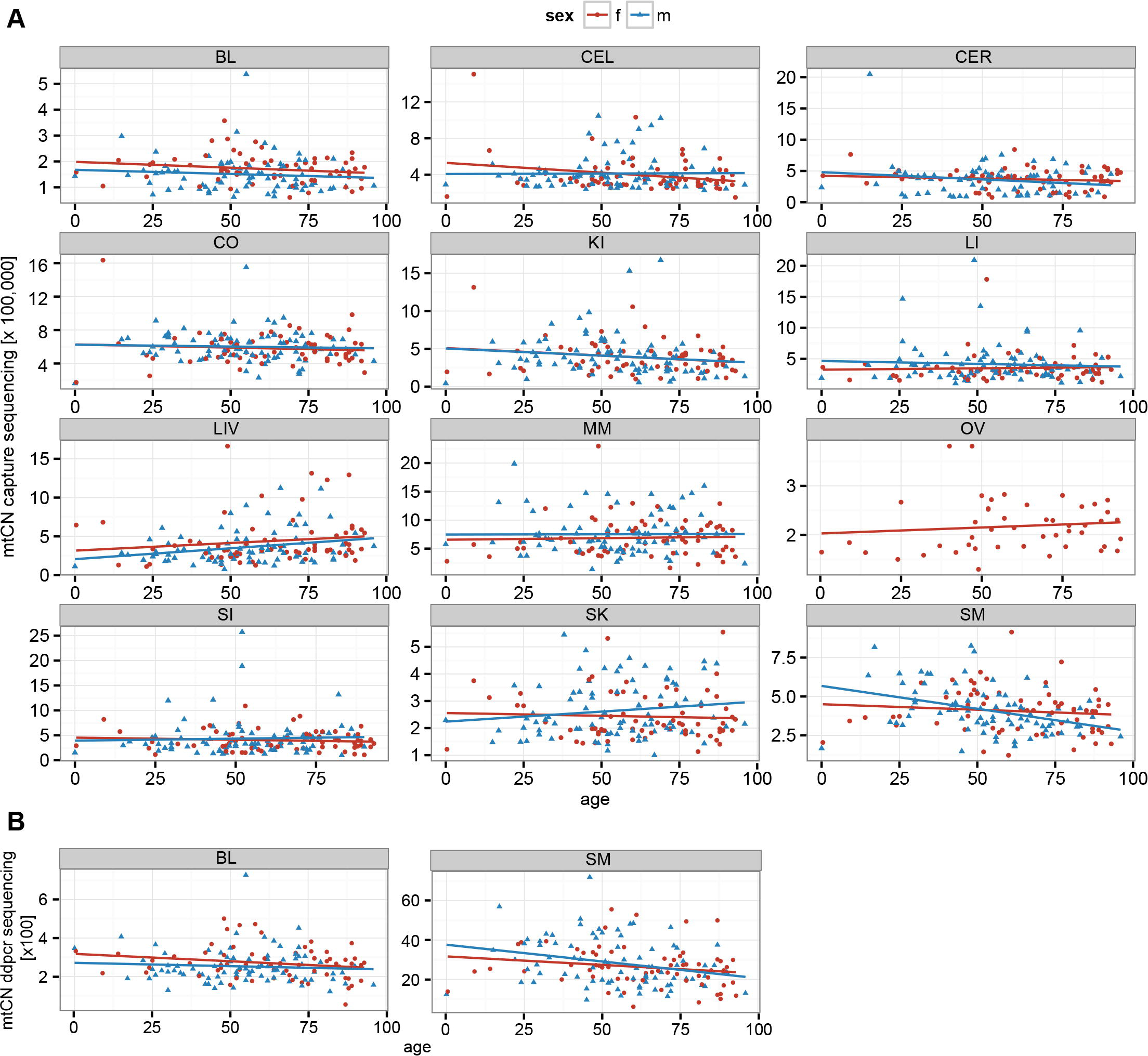
Correlation analysis of estimated mtCN with age. Males (m) and females (f) are distinguished. (A) capture-enrichment. (B) ddPCR.

**S6 Fig.**
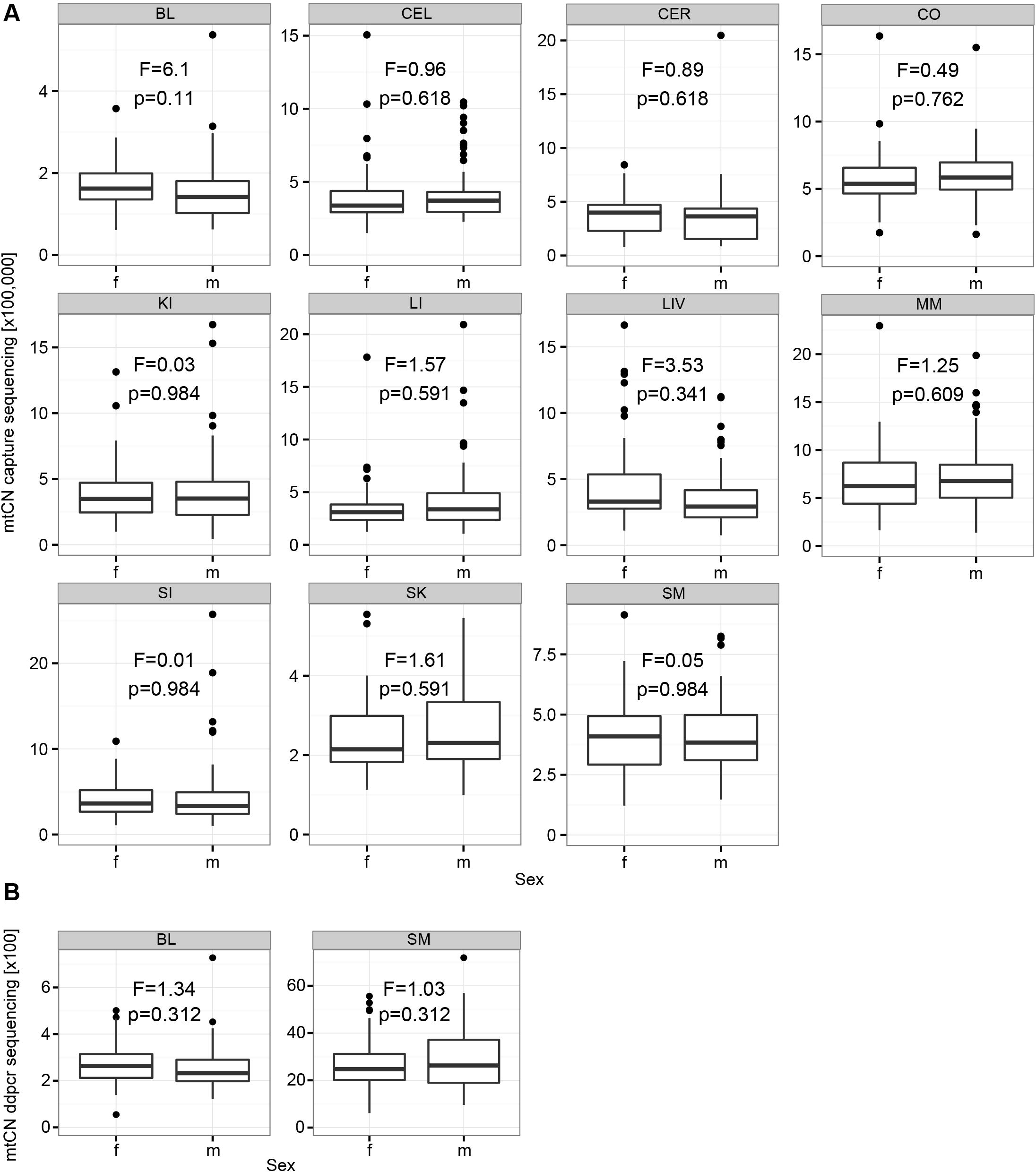
Correlation analysis of mtCN with gender. Males (m) and females (f) are indicated. F- and p-values are specified for each tissue. (A) capture-enrichment. (B) ddPCR.

**S7 Fig.**
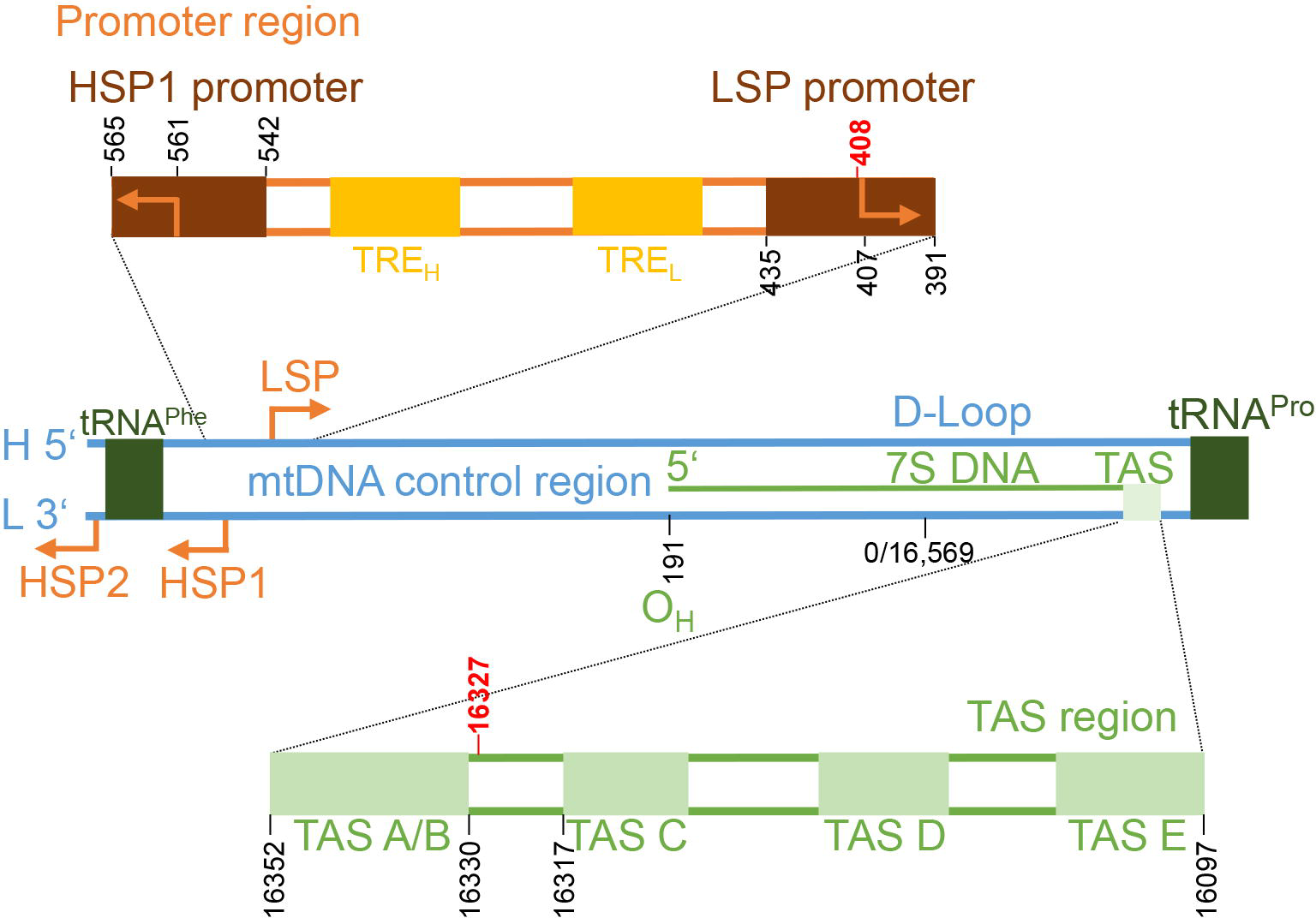
Mitochondrial DNA control region with focus on important regulatory elements for replication. A short RNA primer is transcribed from the light strand promoter (LSP). Replication starts at OH (heavy strand origin of replication). Many replication events terminate in the TAS-region leading to release of a 7S DNA that stays attached to the D-loop region. Positions 408 and 16,327 are located within the LSP or TAS-region, respectively.

**S1 Table. Primers and probes used for ddPCR mtCN estimation.** Sequences are shown from 5’ to 3’. 5’ and 3’ labeling of probes is indicated.

**S2 Table. Correlation analysis of mtCN estimated by different methods with age and sex.** MtCN in the specified tissue was tested for correlation with the indicated test parameter. F-value, correlation coefficient r and p-values (corrected for multiple testing) are given. Tests with significant results (p<0.05) are in red.

**S3 Table. Correlation analysis of mtCN estimates from different methods with haplogroups.** MtCN in samples of the major haplogroups H, J and U were compared to the residual sample set. If mtCNs were determined with different methods, results from all methods are given. F-value, Pearson correlation coefficient r and p-values (corrected for multiple testing) are given. No significant correlations were identified.

**S4 Table. Correlation analysis of mtCN estimates from different methods with the total number of heteroplasmic sites per individual.** The mean and maximum number of heteroplasmic sites per individual are given, along with the F-value, Pearson correlation coefficient r and p-values (corrected for multiple testing). Tests with significant results (p<0.05) are in red.

**S5 Table. Complete list of heteroplasmic sites per tissue investigated for associations between mtCN and MAF.** Only sites that were present in at least 10 individuals in a tissue, colored in red, were tested. The column “Sum” indicates the total number of sites that were tested in a tissue, while the row “Total” indicates the total number of tissues tested for each site.

**S6 Table. Linear regression and Pearson’s correlation analysis of MAF at heteroplasmic sites with mtCN estimates from ddPCR, shotgun sequencing and capture-enrichment sequencing.** Tested sites were found in at least ten individuals in the indicated tissue. F-value, correlation coefficient r and p-values (corrected for multiple testing) are given. Sites with significant correlations with mtCN are marked in red.

**S7 Table. List of all individuals with age, sex, haplogroup, country of origin and mtCN of each tissue estimated with the indicated method.** Samples that were excluded from the data set prior to correlation analysis due to high SD values are marked in red. Empty fields indicates that the sample from this individual was not available. LIV253 was excluded from the data set as inclusion violated the normal distribution of the data set, required for correlation analyses.

## References

1. Stewart JB, Chinnery PF. The dynamics of mitochondrial DNA heteroplasmy: implications for human health and disease. Nat Rev Genet. 2015;16(9):530–42. Epub 2015/08/19. doi: 10.1038/nrg3966. PubMed PMID: 26281784.

2. Shokolenko IN, Alexeyev MF. Mitochondrial DNA: A disposable genome? Biochimica et biophysica acta. 2015;1852(9):1805–9. doi: 10.1016/j.bbadis.2015.05.016. PubMed PMID: MEDLINE:26071375.

3. Montier LLC, Deng JJ, Bai Y. Number matters: control of mammalian mitochondrial DNA copy number. Journal of Genetics and Genomics. 2009;36(3):125–31. doi: 10.1016/s1673-8527(08)60099-5. PubMed PMID: WOS:000264455800001.

4. Miller FJ, Rosenfeldt FL, Zhang CF, Linnane AW, Nagley P. Precise determination of mitochondrial DNA copy number in human skeletal and cardiac muscle by a PCR-based assay: lack of change of copy number with age. Nucleic Acids Research. 2003;31(11). doi: 10.1093/nar/gng060. PubMed PMID: WOS:000183231000003.

5. Liou CW, Lin TK, Chen JB, Tiao MM, Weng SW, Chen SD, et al. Association between a common mitochondrial DNA D-loop polycytosine variant and alteration of mitochondrial copy number in human peripheral blood cells. Journal of Medical Genetics. 2010;47(11):723–8. doi: 10.1136/jmg.2010.077552. PubMed PMID: WOS:000284511700001.

6. Moraes CT. What regulates mitochondrial DNA copy number in animal cells? Trends in Genetics. 2001;17(4):199–205. doi: 10.1016/s0168-9525(01)02238-7. PubMed PMID: WOS:000168718300021.

7. Larsson NG, Wang JM, Wilhelmsson H, Oldfors A, Rustin P, Lewandoski M, et al. Mitochondrial transcription factor A is necessary for mtDNA maintenance and embryogenesis in mice. Nature Genetics. 1998;18(3):231–6. doi: 10.1038/ng0398-231. PubMed PMID: WOS:000072325000020.

8. van Leeuwen N, Beekman M, Deelen J, van den Akker EB, de Craen AJM, Slagboom PE, et al. Low mitochondrial DNA content associates with familial longevity: the Leiden Longevity Study. Age. 2014;36(3):1463–70. doi: 10.1007/s11357-014-9629-0. PubMed PMID: WOS:000342142300037.

9. Ding J, Sidore C, Butler TJ, Wing MK, Qian Y, Meirelles O, et al. Assessing Mitochondrial DNA Variation and Copy Number in Lymphocytes of +2,000 Sardinians Using Tailored Sequencing Analysis Tools. PLoS genetics. 2015;11(7):e1005306-e. doi: 10.1371/journal.pgen.1005306. PubMed PMID: MEDLINE:26172475.

10. Mengel-From J, Thinggaard M, Dalgard C, Kyvik KO, Christensen K, Christiansen L. Mitochondrial DNA copy number in peripheral blood cells declines with age and is associated with general health among elderly. Hum Genet. 2014;133(9):1149–59. Epub 2014/06/07. doi: 10.1007/s00439-014-1458-9. PubMed PMID: 24902542; PubMed Central PMCID: PMCPMC4127366.

11. Frahm T, Mohamed SA, Bruse P, Gemund C, Oehmichen M, Meissner C. Lack of age-related increase of mitochondrial DNA amount in brain, skeletal muscle and human heart. Mechanisms of Ageing and Development. 2005;126(11):1192–200. doi: 10.1016/j.mad.2005.06.008. PubMed PMID: WOS:000232846800008.

12. Bai RK, Perng CL, Hsu CH, Wong LJC. Quantitative PCR analysis of mitochondrial DNA content in patients with mitochondrial disease. In: Lee HK, DiMauro S, Tanaka M, Wei YH, editors. Mitochondrial Pathogenesis: From Genes and Apoptosis to Aging and Disease. Annals of the New York Academy of Sciences. 10112004. p. 304–9.

13. Chu H-T, Hsiao WWL, Tsao TTH, Chang C-M, Liu Y-W, Fan C-C, et al. Quantitative assessment of mitochondrial DNA copies from whole genome sequencing. Bmc Genomics. 2012;13. doi: 10.1186/1471-2164-13-s7-s5. PubMed PMID: WOS:000312987200005.

14. Guo Y, Li J, Li C-I, Shyr Y, Samuels DC. MitoSeek: extracting mitochondria information and performing high-throughput mitochondria sequencing analysis. Bioinformatics. 2013;29(9):1210–1. doi: 10.1093/bioinformatics/btt118. PubMed PMID: WOS:000318573900016.

15. Samuels DC, Li C, Li B, Song Z, Torstenson E, Clay HB, et al. Recurrent Tissue-Specific mtDNA Mutations Are Common in Humans. Plos Genetics. 2013;9(11). doi: 10.1371/journal.pgen.1003929. PubMed PMID: WOS:000330369000022.

16. Michikawa Y, Mazzucchelli F, Bresolin N, Scarlato G, Attardi G. Aging-dependent large accumulation of point mutations in the human mtDNA control region for replication. Science. 1999;286(5440):774–9. doi: 10.1126/science.286.5440.774. PubMed PMID: WOS:000083303200055.

17. Calloway CD, Reynolds RL, Herrin GL, Anderson WW. The frequency of heteroplasmy in the HVII region of mtDNA differs across tissue types and increases with age. American Journal of Human Genetics. 2000;66(4):1384–97. doi: 10.1086/302844. PubMed PMID: WOS:000088373400019.

18. Wang Y, Michikawa Y, Mallidis C, Bai Y, Woodhouse L, Yarasheski KE, et al. Muscle-specific mutations accumulate with aging in critical human mtDNA control sites for replication. Proceedings of the National Academy of Sciences of the United States of America. 2001;98(7):4022–7. doi: 10.1073/pnas.061013598. PubMed PMID: WOS:000167833700075.

19. Li M, Schroeder R, Ni S, Madea B, Stoneking M. Extensive tissue-related and allele-related mtDNA heteroplasmy suggests positive selection for somatic mutations. Proceedings of the National Academy of Sciences of the United States of America. 2015;112(8):2491–6. doi: 10.1073/pnas.1419651112. PubMed PMID: WOS:000349911700055.

20. He Y, Wu J, Dressman DC, Iacobuzio-Donahue C, Markowitz SD, Velculescu VE, et al. Heteroplasmic mitochondrial DNA mutations in normal and tumour cells. Nature. 2010;464(7288):610–U175. doi: 10.1038/nature08802. PubMed PMID: WOS:000275974200052.

21. Naue J, Horer S, Sanger T, Strobl C, Hatzer-Grubwieser P, Parson W, et al. Evidence for frequent and tissue-specific sequence heteroplasmy in human mitochondrial DNA. Mitochondrion. 2015;20:82–94. doi: 10.1016/j.mito.2014.12.002. PubMed PMID: WOS:000348632200011.

22. Avital G, Buchshtav M, Zhidkov I, Tuval J, Dadon S, Rubin E, et al. Mitochondrial DNA heteroplasmy in diabetes and normal adults: role of acquired and inherited mutational patterns in twins. Human Molecular Genetics. 2012;21(19):4214–24. doi: 10.1093/hmg/dds245. PubMed PMID: WOS:000308888100007.

23. Nicholls TJ, Minczuk M. In D-loop: 40 years of mitochondrial 7S DNA. Experimental Gerontology. 2014;56:175–81. doi: 10.1016/j.exger.2014.03.027. PubMed PMID: WOS:000338607100021.

24. Falkenberg M, Larsson N-G, Gustafsson CM. DNA replication and transcription in mammalian mitochondria. Annual Review of Biochemistry. Annual Review of Biochemistry. 762007. p. 679–99.

25. Li H, Durbin R. Fast and accurate short read alignment with Burrows-Wheeler transform. Bioinformatics. 2009;25(14):1754–60. doi: 10.1093/bioinformatics/btp324. PubMed PMID: WOS:000267665900006.

26. Grubbs FE. Sample criteria for testing outlying observations. The Annals of Mathematical Statistics 1950. 27–58 p.

27. Hindson BJ, Ness KD, Masquelier DA, Belgrader P, Heredia NJ, Makarewicz AJ, et al. High-Throughput Droplet Digital PCR System for Absolute Quantitation of DNA Copy Number. Analytical Chemistry. 2011;83(22):8604–10. doi: 10.1021/ac202028g. PubMed PMID: WOS:000296830200035.

28. Li M, Schroeder R, Ko A, Stoneking M. Fidelity of capture-enrichment for mtDNA genome sequencing: influence of NUMTs. Nucleic Acids Research. 2012;40(18). doi: 10.1093/nar/gks499. PubMed PMID: WOS:000309927100001.

29. Renaud G, Stenzel U, Kelso J. leeHom: adaptor trimming and merging for Illumina sequencing reads. Nucleic Acids Research. 2014;42(18). doi: 10.1093/nar/gku699. PubMed PMID: WOS:000347687100005.

30. R Core Team. R: A Language and Environment for Statistical Computing. Vienna, Austria: R Foundation for Statistical Computing; 2015.

31. Benjamini Y, Hochberg Y. Controlling the false discovery rate - a practical and powerful approach to multiple testing. Journal of the Royal Statistical Society Series B-Methodological. 1995;57(1):289–300. PubMed PMID: WOS:A1995QE45300017.

32. Fox J, Weisberg S. An {R} Companion to Applied Regression. Thousand Oaks (California): Sage; 2011.

33. Larsson NG, Oldfors A, Holme E, Clayton DA. Low-levels of mitochondrial transcription factor-A in mitochondrial-DNA depletion. Biochemical and Biophysical Research Communications. 1994;200(3):1374–81. doi: 10.1006/bbrc.1994.1603. PubMed PMID: WOS:A1994NL38800030.

34. Facucho-Oliveira JM, Alderson J, Spikings EC, Egginton S, John JCS. Mitochondrial DNA replication during differentiation of murine embryonic stem cells. Journal of Cell Science. 2007;120(22):4025–34. doi: 10.1242/jcs.016972. PubMed PMID: WOS:000251144700013.

35. Cai N, Chang S, Li Y, Li Q, Hu J, Liang J, et al. Molecular Signatures of Major Depression. Current Biology. 2015;25(9):1146–56. doi: 10.1016/j.cub.2015.03.008. PubMed PMID: WOS:000353999000021.

36. Malik AN, Czajka A. Is mitochondrial DNA content a potential biomarker of mitochondrial dysfunction? Mitochondrion. 2013;13(5):481–92. doi: 10.1016/j.mito.2012.10.011. PubMed PMID: WOS:000323870600012.

37. Wu ZD, Puigserver P, Andersson U, Zhang CY, Adelmant G, Mootha V, et al. Mechanisms controlling mitochondrial biogenesis and respiration through the ι thermogenic coactivator PGC-1. Cell. 1999;98(1):115–24. doi: 10.1016/s0092-18674(00)80611-x. PubMed PMID: WOS:000081451000013.

38. Gallo RL, Hooper LV. Epithelial antimicrobial defence of the skin and intestine. Nature Reviews Immunology. 2012;12(7):503–16. doi: 10.1038/nri3228. PubMed PMID: WOS:000305803000010.

39. Gomez-Duran A, Pacheu-Grau D, Lopez-Gallardo E, Diez-Sanchez C, Montoya J, Lopez-Perez MJ, et al. Unmasking the causes of multifactorial disorders: ι OXPHOS differences between mitochondrial haplogroups. Human Molecular Genetics. 2010;19(17):3343–53. doi: 10.1093/hmg/ddq246. PubMed PMID: WOS:000280704800005.

40. Suissa S, Wang Z, Poole J, Wittkopp S, Feder J, Shutt TE, et al. Ancient mtDNA Genetic Variants Modulate mtDNA Transcription and Replication. Plos Genetics. 2009;5(5). doi: 10.1371/journal.pgen.1000474. PubMed PMID: WOS:000267083000029.

41. Barthelemy C, de Baulny HO, Diaz J, Cheval MA, Frachon P, Romero N, et al. Late-onset mitochondrial DNA depletion: DNA copy number, multiple deletions, and compensation. Annals of Neurology. 2001;49(5):607–17. doi: 10.1002/ana.1002. PubMed PMID: WOS:000168433700009.

42. Schiaffino S, Reggiani C. Fiber types in mammalian skeletal muscles. Physiological Reviews. 2011;91(4):1447–531. doi: 10.1152/physrev.00031.2010. PubMed PMID: WOS:000296561600009.

43. Amara CE, Shankland EG, Jubrias SA, Marcinek DJ, Kushmerick MJ, Conley KE. Mild mitochondrial uncoupling impacts cellular aging in human muscles in vivo. Proceedings of the National Academy of Sciences of the United States of America. 2007;104(3):1057–62. doi: 10.1073/pnas.0610131104. PubMed PMID: WOS:000243761100067.

44. Cai N, Li Y, Chang S, Liang J, Lin C, Zhang X, et al. Genetic Control over mtDNA and Its Relationship to Major Depressive Disorder. Current Biology. 2015;25(24):3170–7. doi: 10.1016/j.cub.2015.10.065. PubMed PMID: WOS:000367233400016.

45. Del Bo R, Bordoni A, Boneschi FM, Crimi M, Sciacco M, Bresolin N, et al. Evidence and age-related distribution of mtDNA D-loop point mutations in skeletal muscle from healthy subjects and mitochondrial patients. Journal of the Neurological Sciences. 2002;202(1-2):85–91. doi: 10.1016/s0022-510x(02)00247-2. PubMed PMID: WOS:000178254900014.

46. Hebert SL, Marquet-de Rouge P, Lanza IR, McCrady-Spitzer SK, Levine JA, Middha S, et al. Mitochondrial Aging and Physical Decline: Insights From Three Generations of Women. Journals of Gerontology Series a-Biological Sciences and Medical Sciences. 2015;70(11):1409–17. doi: 10.1093/gerona/glv086. PubMed PMID: WOS:000364765700015.

47. Annex BH, Williams RS. Mitochondrial-DNA structure and expression in specialized subtypes of mammalian striated-muscle. Molecular and Cellular Biology. 1990;10(11):5671–8. PubMed PMID: WOS:A1990ED48400009.

48. Roberti M, Musicco C, Polosa PL, Milella F, Gadaleta MN, Cantatore P. Multiple protein-binding sites in the TAS-region of human and rat mitochondrial DNA. Biochemical and Biophysical Research Communications. 1998;243(1):36–40. doi: 10.1006/bbrc.1997.8052. PubMed PMID: WOS:000071862400007.

49. Levin L, Mishmar D. A genetic view of the mitochondrial role in ageing: killing us softly. Advances in experimental medicine and biology. 2015;847:89–106. doi: 10.1007/978-1-4939-2404-2_4. PubMed PMID: MEDLINE:25916587.

50. Wallace DC, Chalkia D. Mitochondrial DNA Genetics and the Heteroplasmy Conundrum in Evolution and Disease. Cold Spring Harbor Perspectives in Biology. 2013;5(11). doi: 10.1101/cshperspect.a021220. PubMed PMID: WOS:000327742400011.

51. Rooney AP, Zhang JZ. Rapid evolution of a primate sperm protein: Relaxation of functional constraint or positive Darwinian selection? Molecular Biology and Evolution. 1999;16(5):706–10. PubMed PMID: WOS:000080208300014.

52. Bjornerfeldt S, Webster MT, Vila C. Relaxation of selective constraint on dog mitochondrial DNA following domestication. Genome Research. 2006;16(8):990–4. doi: 10.1101/gr.5117706. PubMed PMID: WOS:000239441400005.

53. Wang XX, Thomas SD, Zhang JZ. Relaxation of selective constraint and loss of function in the evolution of human bitter taste receptor genes. Human Molecular Genetics. 2004;13(21):2671–8. doi: 10.1093/hmg/ddh289. PubMed PMID: WOS:000224703900012.

54. Diaz F, Bayona-Bafaluy MP, Rana M, Mora M, Hao H, Moraes CT. Human mitochondrial DNA with large deletions repopulates organelles faster than full-length genomes under relaxed copy number control. Nucleic Acids Research. 2002;30(21):4626–33. doi: 10.1093/nar/gkf602. PubMed PMID: WOS:000179038100006.

55. Conley KE, Amara CE, Jubrias SA, Marcinek DJ. Mitochondrial function, fibre types and ageing: new insights from human muscle in vivo. Experimental Physiology. 2007;92(2):333–9. doi: 10.1113/expphysiol.2006.034330. PubMed PMID: WOS:000244741000006.

56. deGrey A. A proposed refinement of the mitochondrial free radical theory of aging. Bioessays. 1997;19(2):161–6. doi: 10.1002/bies.950190211. PubMed PMID: WOS:A1997WG37300010.

57. van Oven M, Kayser M. Updated Comprehensive Phylogenetic Tree of Global Human Mitochondrial DNA Variation. Human Mutation. 2009;30(2):E386–E94. doi: 10.1002/humu.20921. PubMed PMID: WOS:000279979200007.

58. Jendrach M, Pohl S, Voth M, Kowald A, Hammerstein P, Bereiter-Hahn J. Morpho-dynamic changes of mitochondria during ageing of human endothelial cells. Mechanisms of Ageing and Development. 2005;126(6-7):813–21. doi: 10.1016/j.mad.2005.03.002. PubMed PMID: WOS:000229536000023.

59. Figge MT, Reichert AS, Meyer-Hermann M, Osiewacz HD. Deceleration of Fusion-Fission Cycles Improves Mitochondrial Quality Control during Aging. Plos Computational Biology. 2012;8(6). doi: 10.1371/journal.pcbi.1002576. PubMed PMID: WOS:000305965300042.

